# Decoding the role of CBLB for innate immune responses regulating systemic dissemination during Non-Tuberculous Mycobacteria infection

**DOI:** 10.1101/2020.05.28.117663

**Authors:** Srinivasu Mudalagiriyappa, Jaishree Sharma, Hazem F. M. Abdelaal, Thomas C. Kelly, Woosuk Choi, Miranda D. Vieson, Adel M. Talaat, Som G. Nanjappa

## Abstract

Non-Tuberculous Mycobacteria (NTM) are ubiquitous in nature, present in soil and water, and cause primary leading to disseminated infections in immunocompromised individuals. NTM infections are surging in recent years due to an increase in an immune-suppressed population, medical interventions, and patients with underlying lung diseases. Host regulators of innate immune responses, frontiers for controlling infections and dissemination, are poorly defined during NTM infections. Here, we describe the role of CBLB, an E3-ubiquitin ligase, for innate immune responses and disease progression in a mouse model of NTM infection under compromised T-cell immunity. We found that CBLB thwarted NTM growth and dissemination in a time- and infection route- dependent manner. Mechanistically, we uncovered defects in many innate immune cells in the absence of *Cblb*, including poor responses of NK cells, inflammatory monocytes, and conventional dendritic cells. Strikingly, Cblb-deficient macrophages were competent to control NTM growth *in vitro*. Histopathology suggested the lack of early formation of granulomatous inflammation in the absence of CBLB. Collectively, CBLB is essential to mount productive innate immune responses and help prevent the dissemination during an NTM infection under T-cell deficiency.

## Introduction

Nontuberculous mycobacteria (NTM) or atypical mycobacteria, distinct from tuberculosis causing mycobacteria, are widely distributed in the environment. However, among 190 species, only a handful of NTM, predominantly of *Mycobacterium avium* complex (MAC) species are isolated from infected patients. Infections caused by MAC are globally prevalent (47%-80%), causing mortality up to 42% (Prevots et al., 2017; Ruth and van Ingen, 2017; Spaulding et al., 2017; Diel et al., 2018). Further, it is estimated that NTM infections are rising at a rate of 8% annually associated with increasing immune-suppressed population, including the patients with underlying lung diseases and the geriatric population (Adjemian et al., 2012; Winthrop et al., 2019). Thus, with the global prevalence and ubiquitous nature of bacteria, NTM infections are increasing in a susceptible population at an alarming rate. The emergence of drug resistance and poor understanding of protective immune correlates of NTM infections, further complicate the disease control and prevention (Horne and Skerrett, 2019). Innate immune cell responses, key orchestrators of initial infection control and adaptive immunity, are poorly defined during NTM infections.

Casitas B-lineage lymphoma (Cblb) is an E3 ubiquitin ligase shown to regulate both innate and adaptive immune responses, with a well-studied role for adaptive immune responses (Liu et al., 2014; Tang et al., 2019). CBLB is widely expressed in peripheral lymphoid cells, including several innate immune cells, and is a well-known negative regulator of T cell responses (Bachmaier et al., 2000; Jeon et al., 2004; Paolino and Penninger, 2010). Ablation of *Cblb* potentiated T cell responses, and *Cblb*-deficient mice spontaneously rejected tumors in a CD8^+^ T-cell dependent manner (Loeser et al., 2007). *Cblb* deficiency facilitated T cell activation independent of CD28 requirement (Chiang et al., 2000), and the lack of CBLB has been implicated in breaking peripheral tolerance and contributing to autoimmunity/allergic responses (Paolino and Penninger, 2010; Oh et al., 2011; Paolino et al., 2011; Lutz-Nicoladoni et al., 2015; Singh et al., 2018; Tang et al., 2019). Further, *Cblb* can be targeted to bolster the CD8^+^ T cell responses for fungal vaccine immunity even in the absence of CD4^+^ T-cell help (Nanjappa et al., 2018), suggesting translational implications. CBLB, in T cells, targets several important signaling pathway proteins, including PLCγ1, VAV1, NEDD4, PKCθ, WASP, and Crk-L, mainly by polyubiquitination and their degradation (Tang et al., 2019). Notably, several SNPs in *Cblb* gene are identified in humans and have been associated with several diseases or disorders that are directly or indirectly implicated with T cells (Kosoy et al., 2004; International Multiple Sclerosis Genetics et al., 2007; Payne et al., 2007; Perez et al., 2010; Sanna et al., 2010; Doniz-Padilla et al., 2011; DeWan et al., 2012; Sturner et al., 2014; Li et al., 2018). Thus, CBLB acts as a negative regulator of T-cell functions by targeting key signaling molecules required for T-cell activation.

While CBLB roles are well defined for adaptive immunity, its functions are poorly described for innate immunity, especially during infections. Accumulating data suggest its diverse functions in many innate immune cells. The development of macrophages, dendritic cells, and NK cells seem to be intact in the absence of *Cblb* (Loeser and Penninger, 2007). However, deficiency of *Cblb* in NK cells potentiated their functions; production of IFNγ and antitumor activity (Yasuda et al., 2002; Paolino et al., 2014; Chirino et al., 2019). In macrophages, CBLB prevented LPS-induced septic shock by downregulating TLR4 (Bachmaier et al., 2007). Recently, CBLB has been shown to downregulate Syk kinase, and its ablation/inhibition led to enhanced production of proinflammatory cytokines, the release of reactive oxygen species (ROS), and fungal killing by macrophages (Wirnsberger et al., 2016; Xiao et al., 2016; Zhu et al., 2016). Further, CBLB has regulatory roles in dendritic cells (DC) by modulating the functions, both positively and negatively (Arron et al., 2001; Wallner et al., 2013; Tang et al., 2019). Nevertheless, CBLB functions in innate immunity during mycobacterial diseases are not deciphered.

In this study, we systematically evaluated the role of CBLB for innate immunity and dissemination of bacteria in a mouse model of NTM infection with deficient T-cell responses. We assessed the bacterial control and innate immune cell numbers/responses following both intratracheal and intravenous infections for up to 5 months. We describe the critical role of CBLB in dictating the pathogenesis of NTM infection that was associated with multiple defective or altered dynamics of innate immune cell numbers or their responses.

## Materials and Methods

### Mice

The OT-I Tg (TCRα/TCRβ specific for OT-I epitope; Stock #: 003831) and B6.PL-Thy1a/Cy/Thy1.1 (Thy1.1; Stock #: 000406) were purchased from Jackson Laboratories. *Cblb*^-/-^ mice were provided by P.S. Ohashi (University of Toronto, Ontario, Canada) with permission from Josef Penninger (IMBA, Austria). OTI-Tg mice were backcrossed with *Cblb*^*-/-*^ to generate OT-I Tg-*Cblb*^-/-^ mice at the UW-Madison facility and were transferred to the current facility with the facilitation from Bruce Klein, UW-Madison. All mice were maintained under specific-pathogen-free conditions at the University of Illinois at Urbana-Champaign (UIUC). All experiments were conducted in accordance with the guidelines of the Institutional Animal Care and Use Committee of the UIUC.

### Infections

*Six- to eight*-week-old mice were used for all the infections in this study. Recombinant dsRED^+^ Kanamycin-resistant *Mycobacterium avium* strain 104 (MAV 104) was cultured in Middlebrook 7H9 broth (Difco) supplemented with albumin, dextrose and catalase (ADC) and Kanamycin, and at ∼OD of 0.8-1.0, culture was centrifuged and resuspended in sterile PBS for infection. Mice were either infected intravenously (I.V.) with 1×10^6^ Colony Forming Units (CFU) or intratracheally (I.T. by intubation under sedation) with 1×10^5^ CFU.

### *In vitro* experiments

For *in vitro* infections, bone-marrow-derived cells (Macrophages-BMM: GM-CSF-10ng/ml for six days) were infected with either 5 MOI or 10 MOI (Multiplicity of Infection; bacteria:cells) of bacteria in RPMI supplemented with 10% FBS (complete media). After 4hr incubation, BMM cells were washed to remove any floating/non-adherent bacteria, and were incubated further in complete media. Bone-marrow-derived neutrophils (BMN), isolated following the protocol (Swamydas and Lionakis, 2013), were infected with 5 MOI of dsRED^+ve^ MAV104 and analyzed by flow cytometry.

### Bacterial burden in tissues/BMM cultures

Infected cells or tissues were harvested, homogenized, and plated on BD Middlebrook 7H10 agar plates supplemented with ADC and Kanamycin. Infected BMMs were lysed using 1% Triton X-100 before plating.

### Flow Cytometry

The tissues (lung and spleen) were harvested on indicated time-points, single-cell suspensions were prepared using BD Cell Strainers, and RBCs were lysed using 4% ammonium chloride containing buffer. Similarly, bone-marrow-derived cells were harvested from the plates either by scraping (BMM) or collecting (BMN) at indicated times during *in vitro* experiments. Cells were then stained with the fluorochrome-conjugated antibodies (BD Biosciences, Biolegend, and Invitrogen) along with Live/Dead staining (Invitrogen) for 30’ at 4° C in the dark. For measuring cytokine production by NK cells, infected cells were incubated with Golgi Stop (BD Biosciences) for the last 4hrs before subjecting for fluorochrome-conjugated antibody staining for surface markers and intracellular cytokine (Perm/Fix buffer, BD Biosciences). Cells were analyzed by 24-color compatible *full-spectrum* Cytek Aurora flow analyzer (College of Veterinary Medicine, UIUC).

### Confocal Microscopy

BMM cells were plated on micro-slide glass bottom wells (ibidi) a day before infection. At 48-hr post-infection, wells were washed with PBS, and stained with dyes (DAPI, Lysotracker, and CellROX; Molecular Probes) for 30’ at 37° C. Wells were washed and resuspended in complete media before the microscopy. Images (at least ten distinct fields) were taken on 4-laser Nikon A1R confocal microscope (College of Veterinary Medicine, UIUC) at 60x plan apo λ/1.40 oil, and analyzed using NIS-Elements C software. For quantifying the ROS production in BMM cells from confocal images, ImageJ software of the National Institute of Health (http://rsweb.nih.gov.ij/) was used for each cell of the representative field.

### Histopathological studies

Tissues were harvested and stored in 10% Buffered Formalin containers. Tissues were paraffin-embedded and sections were mounted on slides for H&E staining. Some sections were also stained for acid-fast bacilli with Ziehl-Neelsen stain. Histopathology was then analyzed and interpreted by a board-certified veterinary anatomic pathologist in a blinded manner.

### Measuring CBLB/Cblb expression

Cells/tissues were harvested and subjected for measuring Cblb RNA and protein by qPCR, Western Blotting, and flow cytometry. For qPCR analysis: The BMM and spleens were harvested, and RNA was extracted using Qiagen RNeasy Kit according to the manufacturer’s instructions. cDNA was synthesized using GoScript Reverse Transcription system (Promega), and qPCR was executed using QuantiNova SYBR Green PCR Kit (Qiagen) by Quant Studio 3 (Applied Biosystems) analyzer. Sequences of primers used are: Cblb-For: CACCCTTCTCCCAAGCATAA, Rev: AGACCGAACAGGAGCTTTGA; β-actin-For: TGGAGAAGAGCTATGAGCTGCCTG, Rev: GTGCCACCAGACAGCACTGTGTTG; and CCL2-For: GAAGGAATGGGTCCAGACAT, Rev: ACGGGTCAACTTCACATTCA. Western Blotting: Cells from spleen and BMM were washed 3 times with ice-cold PBS and lysed by M-PER™ Mammalian Protein Extraction Reagent (Thermo Scientific) in the presence of protease inhibitor cocktail (Sigma-Aldrich) and phosphatase inhibitor cocktail (Sigma-Aldrich). Quantity of cell lysate protein was measured by Pierce™ BCA protein assay kit. CBLB and GAPDH in the protein blots were probed with anti-CBLB- and anti-GAPDH-antibodies (SC-8006 and SC-166545, respectively; Santa Cruz Biotechnology). Flow Cytometry analysis: Cells were stained with surface markers followed by intracellular staining for CBLB (anti-CBLB antibody, G-1, Santa Cruz Biotechnology), and the levels were analyzed by Cytek Aurora analyzer.

### Statistical Analysis

All statistical analyses were performed using a two-tailed unpaired Student t-test using GraphPad Prism 8 software. A two-tailed P value of ≤0.05 was considered statistically significant.

## Results

### Induction of CBLB following NTM infection *in vitro*

CBLB expression is dynamically regulated in T- and myeloid-cells. Following activation of T cells with CD28 stimulation, CBLB expression is downregulated, whereas engaging with CTLA4 and PD-1 receptors upregulate CBLB levels (Li et al., 2004; Karwacz et al., 2011). On a similar note, CBLB was upregulated in the macrophages and dendritic cells following fungal infection (Wirnsberger et al., 2016). Therefore, we were interested in assessing the levels of CBLB following NTM infection. First, we measured the basal levels of CBLB expression in macrophages using bone-marrow-derived macrophages (BMM) and primary cells (peritoneal macrophages). We found detectable levels of CBLB in uninfected *Cblb*^*+/+*^ BMM (**Fig. 1A**; left panel) and was absent in *Cblb*^*-/-*^ cells. Similarly, we measured CBLB levels in peritoneal macrophage cells (good source of primary macrophages; CD11b^+^F4/80^+^) and found appreciable basal levels of CBLB in *Cblb*^*+/+*^ cells (**Fig. 1A**; middle panel). We found a similar phenotype with peritoneal dendritic cells (CD11c^+^F4/80^-^; **Fig. 1A**; right panel). Next, we measured the induction of *Cblb* (mRNA levels) following an NTM (*Mycobacterium avium 104*; MAV104) infection, and found that levels of *Cblb* were reduced in *Cblb*^*+/+*^ BMM (**Fig. 1B**; left panel; protein levels were undetectable by Western blotting-not shown). On the contrary, mRNA levels of *Cblb* were increased in splenocytes after infection (**Fig. 1B**; right panel). Next, we measured the protein levels of CBLB in BMM, splenocytes, and peritoneal macrophages following the infection, and found that CBLB levels were significantly higher in infected *Cblb*^*+/+*^ cells compared with *Cblb*^*-/-*^ cells (**Fig. 1C & D**). Collectively, our data indicated that NTM infection enhances the levels of CBLB in myeloid cells.

**Figure 1:**
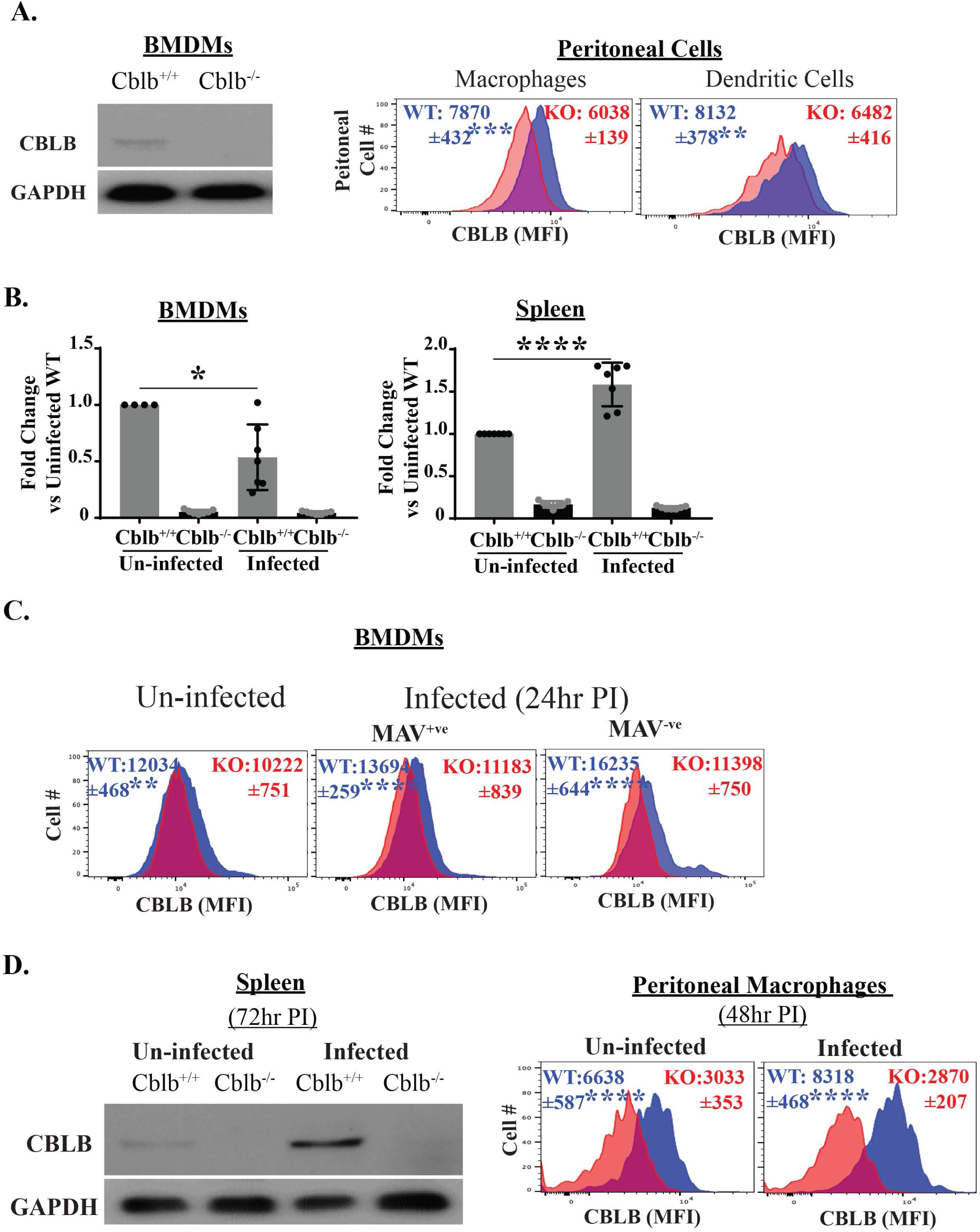
Cblb induction and quantification. mRNA/protein levels of CBLB was measured in naïve or infected cells & tissues by western blotting (**A**; uninfected **& D**; 5 MOI; ∼1×10^6^ cells), flow cytometry (**A**; uninfected, **C and D**; 5 MOI, >1×10^5^ cells @24-48hr post-infection, PI) and qPCR (**B**, pooled data from 2 experiments @24hr PI, 5MOI, >1×10^6^ cells). Data is representative of at least two independent experiments. N=4-6 replicates for qPCR, and 4-5 mice for flow cytometry. Pooled samples were used for western blotting. Values are mean ± SD. *p≤0.05, **p≤0.01, ***p≤0.001, and ****p≤0.0001. MFI=Mean Fluorescence Intensity. The cells/tissues were from both OT-I Tg/KO or WT/KO background mice.

### Ablation of *Cblb* promotes NTM growth and dissemination under T-cell deficiency in mice

Evidence of the role of CBLB for innate immunity during infections is scarce. Recent reports suggest that CBLB constrains innate immune responses during fungal infections, and the loss of CBLB or functions enhances antifungal activities of DC & macrophages to bolster protection from lethal infection (Wirnsberger et al., 2016; Xiao et al., 2016; Zhu et al., 2016; Nanjappa et al., 2018). To investigate the role of CBLB in innate immune cells during an NTM infection, we used OT-I Tg mice (CD8^+^ T-cell transgenic mice, where all T cells are specific for OT-I epitope of ovalbumin) and OT-I-Tg-*Cblb* KO (OT-I Tg lacking Cblb) as *Cblb*^*+/+*^ and *Cblb*^*-/-*^, respectively (in this study). We used these mice for three main reasons: 1. Adaptive T-cell immunity is severely compromised as most of the T cells are CD8^+^ T cells and do not recognize NTM, and have a very low number of CD4^+^ T cells; 2. Lymphoid architecture is not affected as seen with *Rag*^*-/-*^ mice (Koning and Mebius, 2012); and 3. NTM infections commonly occur in individuals with compromised T-cell functions (Henkle and Winthrop, 2015; Ratnatunga et al., 2020). We used a strain of *Mycobacterium avium* (MAV) as an NTM for our studies as most of the NTM infections (50-90%) seen in humans, accounting for up to ∼40% mortality, are caused by M. avium complex (MAC) species (Johnson and Odell, 2014; Diel et al., 2018; Horne and Skerrett, 2019). Mice were infected by both intravenous (i.v.) and intratracheal (i.t.) routes and rested for several weeks to determine the bacterial load in the tissues and dissemination. We found that *i.v.* infection of *Cblb*^*+/+*^ mice caused a gradual increase of bacterial burden in the lungs from Wk 6 to 22 post-infection (PI). In the liver and spleen, NTM bacterial kinetics was either stable or increased during the later time points (**Fig. 2A**). Similarly, following intratracheal infection, the bacterial burden was increased steadily in the lungs and spleens, including mediastinal lymph nodes (draining LN of the lungs), in *Cblb*^*+/+*^ mice (**Fig. 2B**). In striking contrast, the bacterial loads were significantly higher in *Cblb*^*-/-*^ mice compared with *Cblb*^*+/+*^ group at all the timepoints following *i.v.* infection (**Fig. 2A**). Analogously, *Cblb*^-/-^ mice had significantly higher bacterial loads in most tissues compared with *Cblb*^*+/+*^, following *i.t.* infection (**Fig. 2B**). Further, we determined the bacterial load in the brains and found that *Cblb*^-/-^ mice had relatively higher CFUs compared with *Cblb*^+/+^ groups (**Supp. Fig. 1**). Collectively, our data suggest that CBLB inhibits the bacterial growth and dissemination during NTM infection under T-cell deficiency.

**Figure 2:**
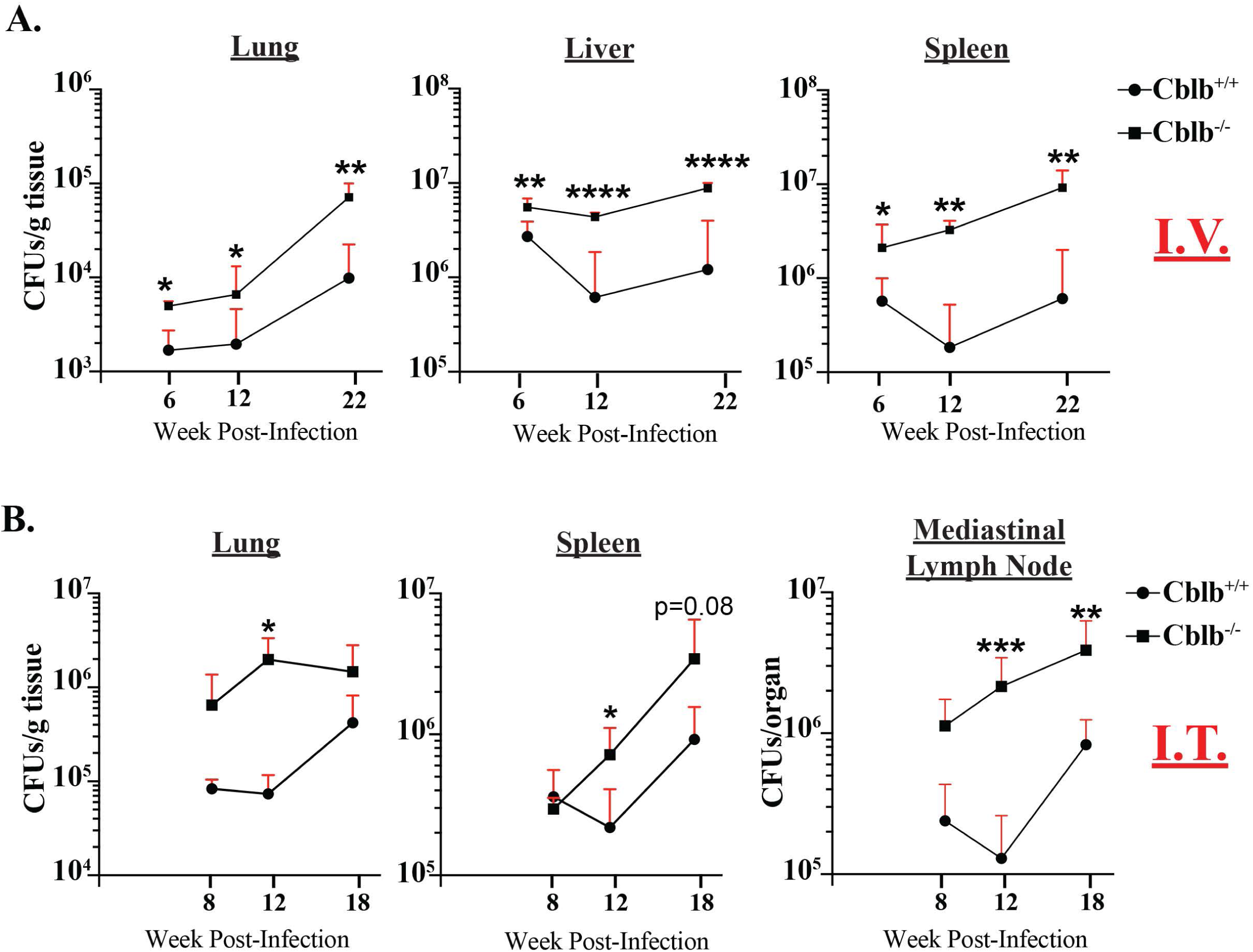
*Cblb*-deficiency promotes NTM dissemination in mice. *Six to eight-*wk-old *OTI-Tg-Cblb*^*+/+*^ (*Cblb*^*+/+*^) and *OTI-Tg-Cblb*^*-/-*^ (*Cblb*^*-/-*^) mice were infected either intravenously (I.V.; **A**) or intratracheally (I.T.; **B**) as described in Methods. At indicated week PI, tissues were harvested and bacterial loads were quantified. IV infection data is representative of 2-3 experiments. Values are mean ± SD. N=3-6 mice/group. *p≤0.05, **p≤0.01, ***p≤0.001, and ****p≤0.0001.

### *Cblb*-deficient macrophages constrain NTM growth *in vitro*

Our *in vivo* bacterial burden and dissemination data (**Fig. 2**) immediately piqued our interest in dissecting the cell-intrinsic role of CBLB in macrophages, the primary target cells of mycobacteria. We infected the bone-marrow-derived macrophages (BMM) *in vitro* to determine bacterial growth (**Fig. 3A-C**). Following 4hr post-infection, BMM were washed to remove any extracellular or non-adherent bacteria and were further incubated for several days. By day 2, with 5 MOI, *Cblb*^-/-^ macrophages controlled the NTM growth significantly better than *Cblb*^*+/+*^ BMM, the phenotype maintained through day 6 (**Fig. 3A**). Although we did not notice significant differences in the later time points (**Fig. 3B**), *Cblb*^*-/-*^ BMM consistently had a lower bacterial burden compared with *Cblb*^*+/+*^ BMM. Further, we tested with a higher infection dose (10 MOI), and the data recapitulated the lower MOI results in that *Cblb*^*-/-*^ BMM controlled NTM growth significantly better, if not completely, than *Cblb*^*+/+*^ BMM (**Fig. 3C**). Further, we tested if it is due to differences in the rate of phagocytosis, and found a similar intake of bacteria by both *Cblb*^*+/+*^ and *Cblb*^*-/-*^ BMM (**Fig. 3D**). To evaluate possible underlying factors for enhanced functions of *Cblb*-deficient BMM, we assessed the activation status and ROS production. We found, following infection, a significant augmented level of MHC-II (MFIs by flow cytometry) in *Cblb*^*-/-*^ BMM compared with *Cblb*^*+/+*^ (**Fig. 3E**). In a similar note, we observed CBLB diminished the production of ROS, if not significantly (**Fig. 3F**). In another approach, we infected *Cblb*^*+/+*^ and *Cblb*^*-/-*^ BMM with dsRED^+^ MAV104, and at 48hr, cells were stained with dyes for confocal microscopy imaging and quantification (**Fig. 3G & H**). We found similar or enhanced staining of CellROX (Invitrogen; measuring cellular oxidative stress) that was associated with lysosomes (Lysotracker; Invitrogen) in both uninfected *Cblb*^*+/+*^ and *Cblb*^*-/-*^ BMM (**Fig. 3G, top panels, and 3H**). In contrast, infected *Cblb*^*+/+*^ BMM had significantly less staining for CellROX compared with *Cblb*^*-/-*^ cells (**Fig. 3G, bottom panels, and 3H**), suggesting a negative role of CBLB for CellROX production. Thus, our data suggest that *Cblb*-deficient macrophages are *competent* in controlling NTM growth.

**Figure 3:**
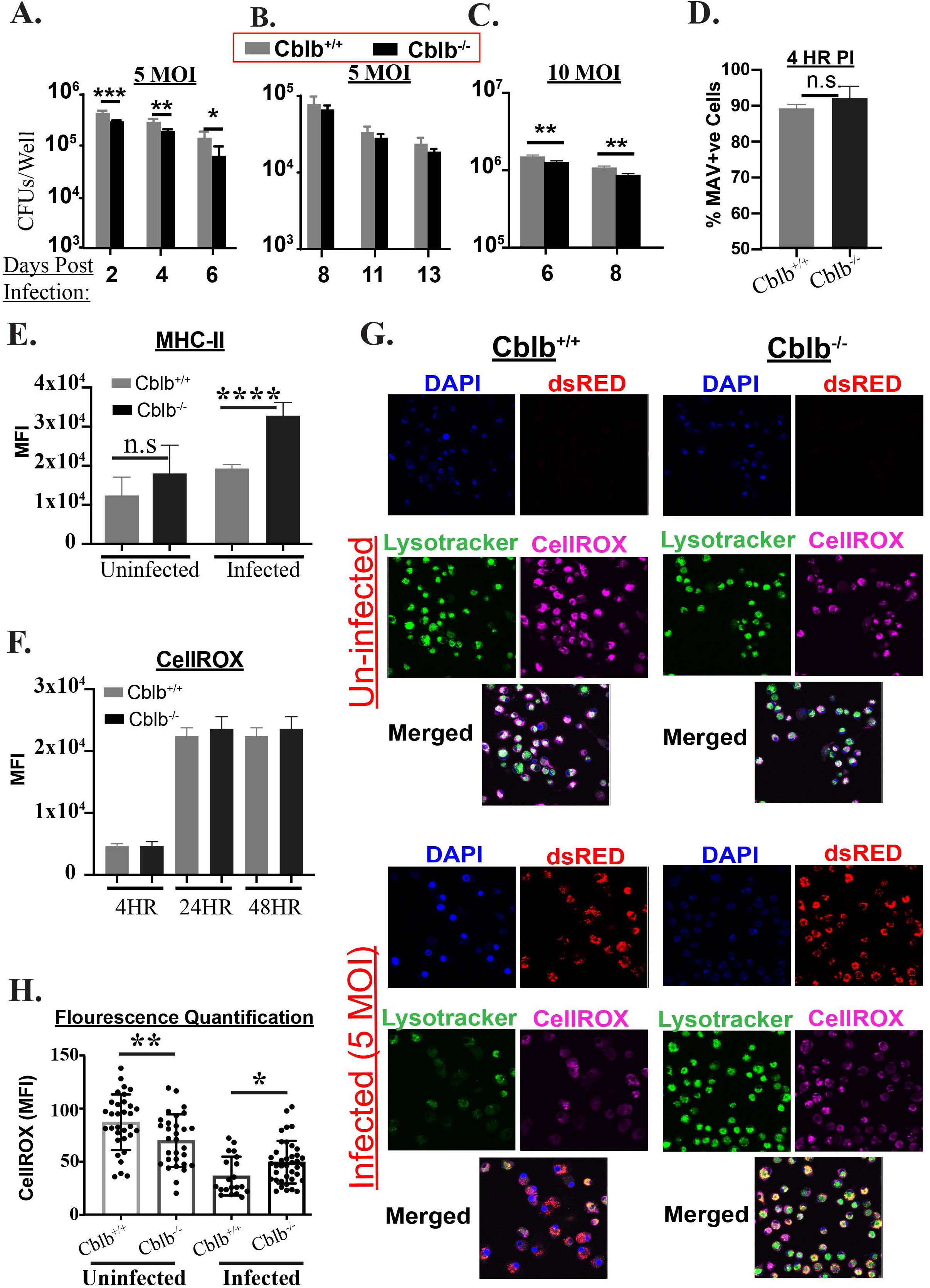
CBLB hinders NTM growth in Bone-Marrow Derived Macrophages (BMM) *in vitro*. Plated BMM cells, either from Tg or WT –background, were infected with MAV104, and at 4hr post-infection, wells were washed to remove non-adherent/extracellular bacteria. On indicated post-infection days, seeded cells (**A**, ∼1×10^6^ cells; **B**, ∼3.5×10^5^ cells; & **C**, 2×10^5^ cells) were lysed with 1% Triton X-100 and CFUs were enumerated on 7H10 agar plates. Data is of 4-replicates/experiment from at least two-independent experiments. After 4hr post-infection, rate of phagocytosis (% dsRED^+ve^; **D**), MHC-II (**E**), and CellROX (**F**) expression in BMM cells were analyzed by flow cytometry. (**G**) Plated infected BMM were washed after 48hr, stained with dyes, and analyzed by confocal microscopy. The ROS levels (**H**) were quantified using Image J software (NIH). Images are representative of 10-fields/experiment of 2-3 independent experiments.

### Flow cytometric analysis of innate immune cells during an NTM infection

Next, we wanted to systematically analyze the dynamics of various innate immune cell numbers/responses during the infection *in vivo* to uncover the possible mechanisms at cellular levels to decipher the higher bacterial growth/dissemination in *Cblb*^*-/-*^ mice. We used several fluorochrome-conjugated antibodies against different surface markers to phenotype innate immune cells and their activation status for analysis by flow cytometry (Misharin et al., 2013; Hey et al., 2017). The gating strategy for identifying the particular innate immune cell subset is shown in **Fig. 4A & B**. We excluded CD90^+^ cells to purge thymic derived T cell population. Gating for Neutrophils and NK cells were common in both lungs and spleens. However, spleens and lungs have different DC and macrophage subsets and are depicted in **Fig. 4A & B**. In essence, the markers for delineating various innate immune cells are defined as below. *<u>Neutrophils</u>*: CD90^-^, CD11b^+^, Ly6G^+^; *<u>NK cells</u>*: CD90^-^, Ly6G^-^, CD11b^-/+^, NK1.1^+^; *<u>Alveolar Macrophages (lung)</u>*: CD90^-^, Ly6G^-^, NK1.1^-^, CD11b^-^, CD11c^+^, Siglec-F^+^; *CD103*^*+*^ *DCs (lung)*: CD90^-^, Ly6G^-^, NK1.1^-^, CD11b^-^, CD11c^+^, CD103^+^; *<u>Interstitial Macrophages (lung)</u>*: CD90^-^, Ly6G^-^, NK1.1^-^, CD11c^+^, CD11b^+^, MHC-II^+^, CD64^+^; *CD11b*^*+*^ *DC (lung)*: CD90^-^, Ly6G^-^, NK1.1^-^, CD11c^+^, CD11b^+^, MHC-II^+^, CD64^-^; *<u>Eosinophils</u>*: CD90^-^, Ly6G^-^, NK1.1^-^, CD11c^-^, CD11b^+^, Siglec-F^+^; *<u>Inflammatory monocytes</u>*: CD90^-^, Ly6G^-^, NK1.1^-^, CD11c^-^, CD11b^+^, Siglec-F^-^, Ly6C^hi^; *<u>Plasmacytoid DC</u>*: CD90^-^, Ly6G^-^, NK1.1^-^, CD11c^-^, CD11b^-^, Ly6C^+^; *CD11b*^*-*^*/CD8*^*+*^ *DC (spleen)*: CD90^-^, Ly6G^-^, NK1.1^-^, CD11b^-^, MHC-II^+^; *CD11b*^*+*^*/CD8*^*-*^ *DC (spleen)*: CD90^-^, Ly6G^-^, NK1.1^-^, CD11b^+^, MHC-II^+^; *Ly6C*^*+*^; and *<u>Resident Monocytes (spleen)</u>*: CD90^-^, Ly6G^-^, NK1.1^-^, CD11c^-^, CD11b^+^, Ly6C^+^. We used this gating strategy for obtaining all flow cytometric data of innate immune cell numbers/responses during *in vivo* studies.

**Figure 4:**
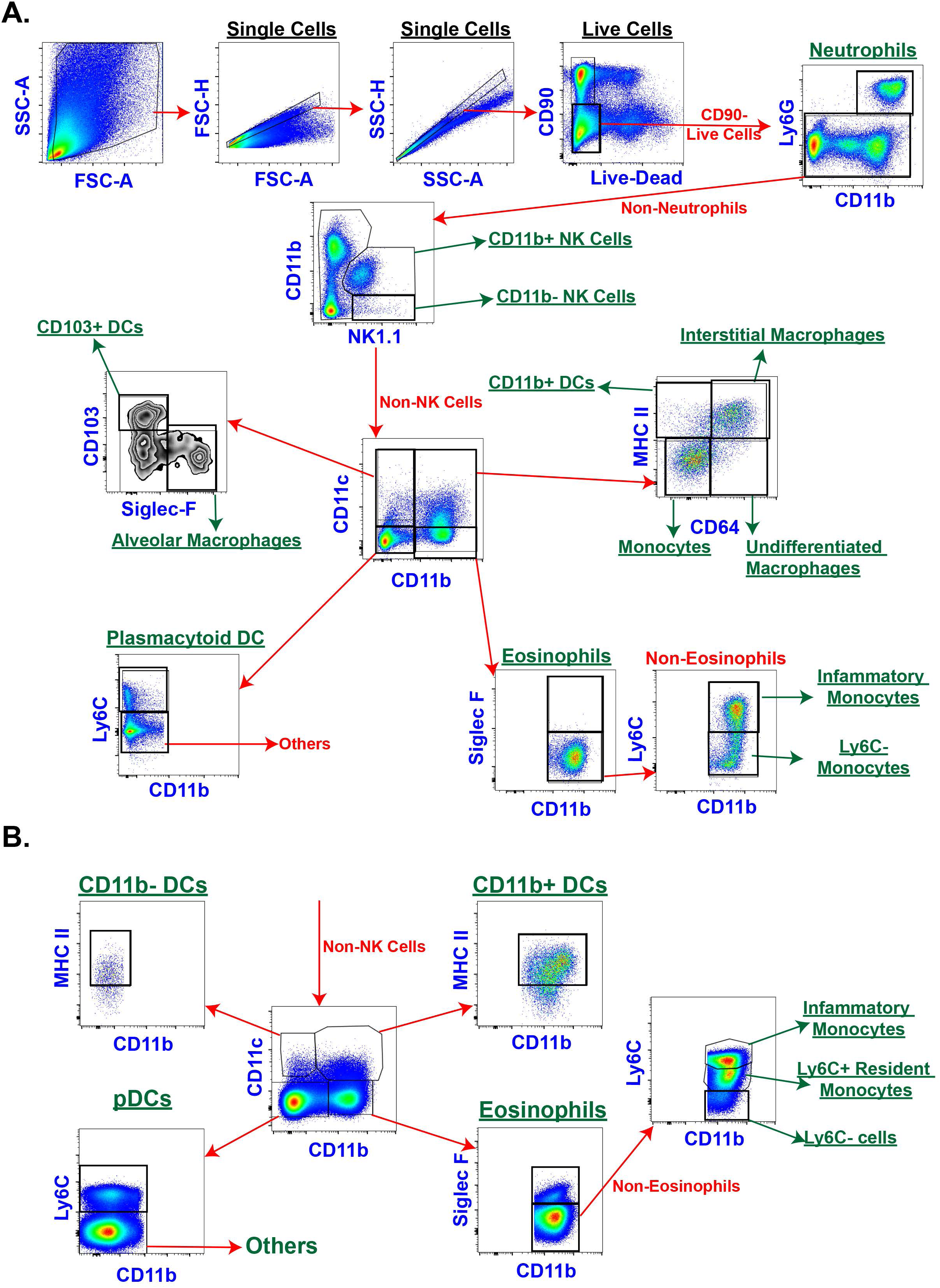
Gating strategy to analyze various innate immune cells by flow cytometry. Mice were infected as described in Fig. 2. Single-cell suspensions from the lung and spleens were stained with fluorochrome-conjugated antibodies and were analyzed by flow cytometry. The figure shows the gating strategy to analyze various innate immune cells and the derivatives for calculation of their frequencies in the lung (**A**) and spleen (**B**; after NK cell gating). The frequencies calculated were on among their immediate parent populations in following figures.

### Deficiency of NK cell responses in the absence of *Cblb* during an NTM infection

We first evaluated NK cell, an important innate immune subset, responses for mediating resistance to mycobacterial infections, including NTM infections (Allen et al., 2015; Garand et al., 2018; Lai et al., 2018a). NK cells are cytotoxic cells, and one of the major IFNγ producing cells protecting against mycobacterial infections. Additionally, it has been shown that CD11b expressing NK cells are more potent in their functions (Fu et al., 2014; Allen et al., 2015; Venkatasubramanian et al., 2017; Cong and Wei, 2019). First, we examined, *in vitro*, if CBLB is expressed in and affected the NK cell functions after infection. The data showed that CBLB expression levels were significantly increased in NK cells following NTM infection (**Fig. 5A**). In congruence with published reports on negative regulation of CBLB in NK cells, we found a significantly higher production of IFNγ in *Cblb*^*-/-*^ cells, irrespective of CD11b subsets, after infection (**Fig. 5B**). Thus, we wanted to enumerate the numbers of CD11b^+^ and CD11b^-^ NK cells during an NTM infection *in vivo*. **Fig. 5C** shows the frequencies of CD11b^+^ NK cells in the lung (left panels) and spleens (right panels) at Wk22 PI following *i.v.* infection. The CD11b^+^ NK cell numbers were significantly lower in *Cblb*^-/-^ compared with *Cblb*^+/+^. The frequencies of NK cells were lower (lung) or significantly reduced (spleen) following *i.v.* infection (**Fig. 5D**). We observed a similar defect, except at Wk 12PI, in CD11b^+^ NK subset numbers following *i.t.* infection (**Fig. 5E**), suggesting that *Cblb*-deficiency blunted CD11b^+^ NK cell responses during an NTM infection. However, we found a non-apparent (i.v. infection) or a significant (i.t. infection) effect of CBLB on CD11b^-^ cell numbers (**Supp. Fig. 2A & B**). Thus, CBLB was required to sustain or potentiate NK cell numbers/responses during an NTM infection.

**Figure 5:**
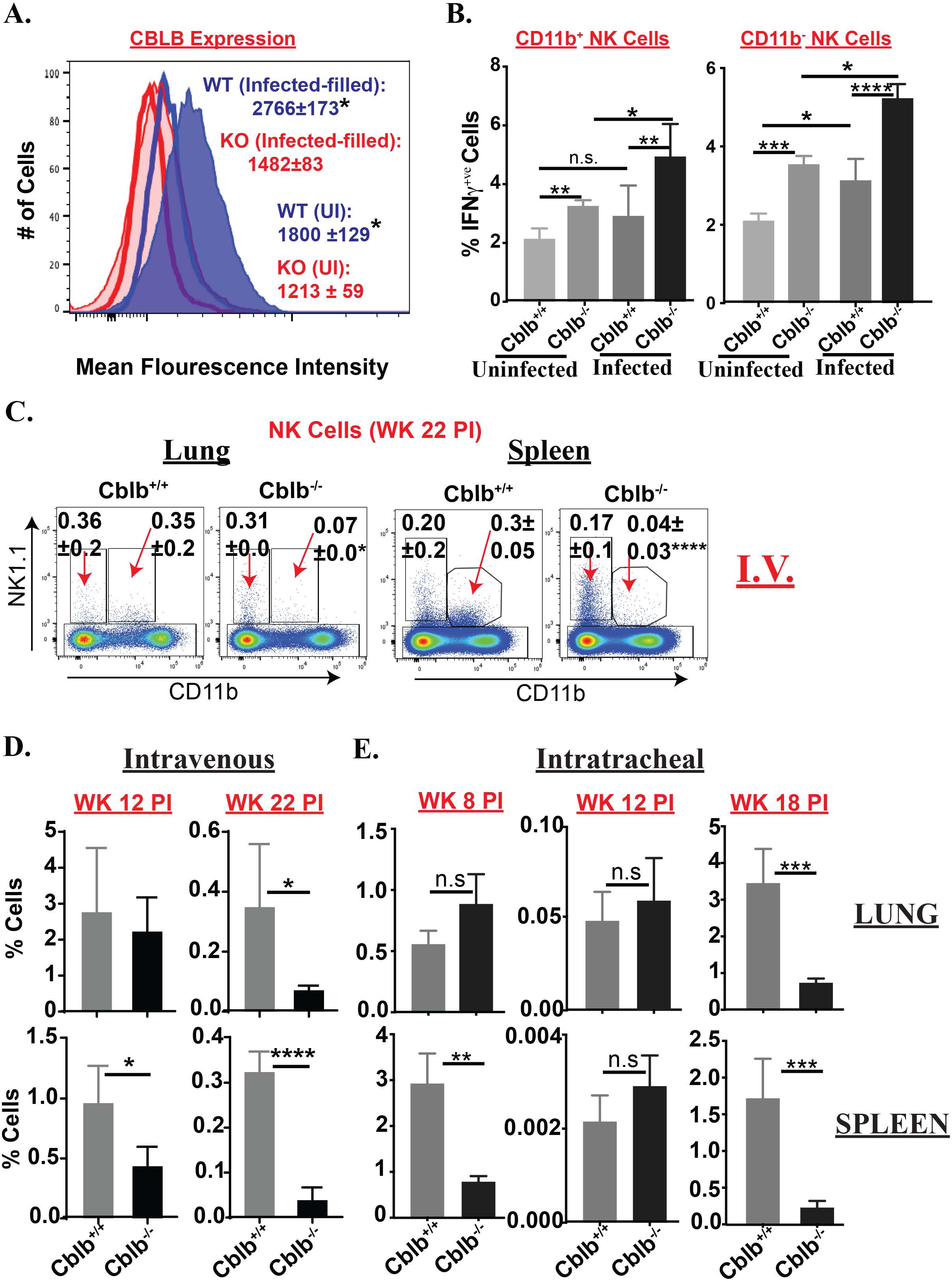
NK cell responses in the absence of *Cblb*. **A & B**. Splenocytes from both OT-I Tg/KO or WT/KO background mice were infected with 5 MOI of MAV104. At 48hr PI, CBLB induction and % cytokine production by NK cells were analyzed by flow cytometry (N=4-6 replicates and data is representative of 3 independent experiments). **C-E**. *Cblb*^*+/+*^and *Cblb*^*-/-*^ mice were infected as described in Fig. 2. At indicated week PI, tissues were harvested, single-cell suspensions were stained for NK cells (CD90^-^, CD11b^+/-^ NK1.1^+^) using fluorochrome-conjugated antibodies, and analyzed by flow cytometry. **C.** Dot plots show the percent of cells. **D & E.** Bar diagrams show the frequencies of NK cells. IV infection data is representative of 2-3 independent experiments. Values are mean ± SD. N=3-6 mice/group. *p≤0.05, **p≤0.01, ***p≤0.001, and ****p≤0.0001.

### Dynamics of alveolar macrophages in the absence of *Cblb* during an NTM infection

Next, we investigated if CBLB regulates the number of alveolar macrophages, implicated in orchestrating mycobacterial diseases (Russell et al., 2009; Cohen et al., 2018). Here, we assessed the alveolar macrophage numbers during NTM infection. Following *i.v.* infection, we found dampened alveolar macrophage numbers in the lungs of *Cblb*^*-/-*^ mice compared with *Cblb*^*+/+*^ group (**Fig. 6A & B**) with a significant reduction during earlier time points, wks 6 & 12 PI. Interestingly, we found no difference of alveolar macrophage numbers in *Cblb* -sufficient and - deficient groups, except at Wk 12, following *i.t.* infection (**Fig. 6C**), suggesting the dichotomous role of CBLB depending on the route of NTM infection.

**Figure 6:**
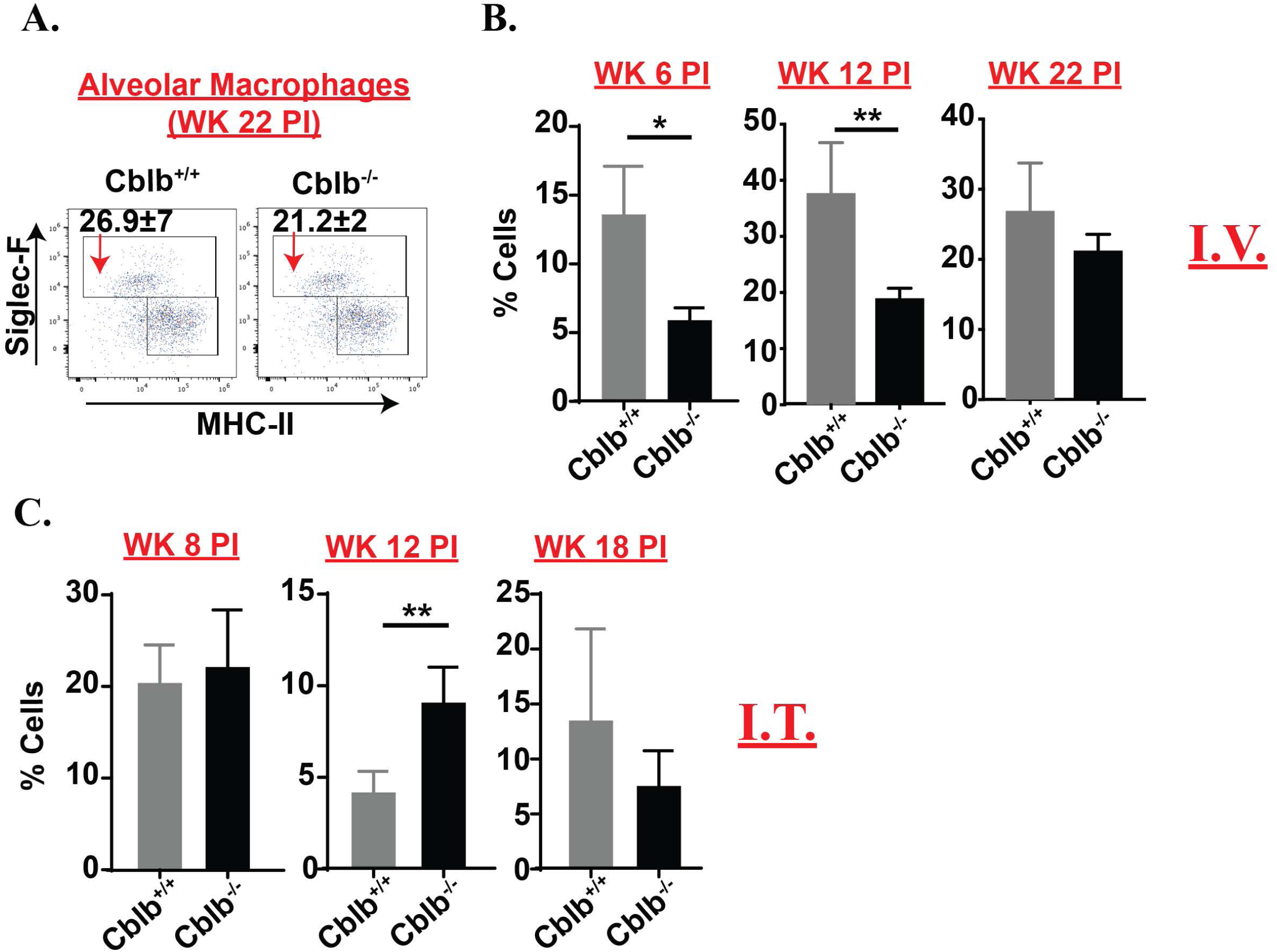
Alveolar Macrophage numbers in the absence of *Cblb*. Mice were infected as described in Fig. 2. At indicated week PI, tissues were harvested, single-cell suspensions were stained for alveolar macrophages (CD90^-^, Ly6G^-^, NK1.1^-^, CD11b^-^, CD11c^+^, Siglec-F/CD64^+^) using fluorochrome-conjugated antibodies, and analyzed by flow cytometry. **A.** Dot plots show percent Alveolar Macrophages. **B & C.** Bar diagrams show the frequencies of Alveolar Macrophages in the lung of mice infected by I.V. and I.T. routes, respectively. Values are mean ± SD. N=3-6 mice/group. IV infection data is representative of 2-3 independent experiments. *p≤0.05 and **p≤0.01.

### *Cblb* deficiency affects inflammatory monocyte responses during an NTM infection

Inflammatory monocytes dictate the outcome of several infections including mycobacterial (Shi and Pamer, 2011; Grainger et al., 2013; Liu et al., 2017; Dunlap et al., 2018; Sampath et al., 2018; Heung and Hohl, 2019), and are part of diverse populations of myeloid cells (Srivastava et al., 2014) triggered during tuberculosis for pathogenesis (Behar et al., 2010; Dorhoi et al., 2014; Lastrucci et al., 2015). Here, we evaluated the role of CBLB for inflammatory monocyte responses during NTM infection. Following *i.v.* infection, sequestration of inflammatory monocytes was overall similar between *Cblb*^*+/+*^ and *Cblb*^*-/-*^ mice in the lungs, but not in the spleen (**Fig. 7A & D**). In the spleen, infected *Cblb*^*-/-*^ mice had significantly lower inflammatory monocytes compared with *Cblb*^*+/+*^ group, and a reciprocal increase in tissue-resident monocytes. Following *i.t.* infection, inflammatory monocytes in both lung and spleen were similar between both *Cblb*^*+/+*^ and *Cblb*^*-/-*^ groups, except at Wk 18 PI, where we found an increase in *Cblb*^-/-^ mice (**Fig. 7D**). Next, we looked at the expression level of MHC-II (as a measure of activation. The data suggested that MHC-II levels, reflected by MFI, were significantly lower on inflammatory monocytes at both Wks 6 and 22 PI in *Cblb*^*-/-*^ groups compared with *Cblb*^*+/+*^ controls (**Fig. 7B**) in both lung and spleens. Additionally, we measured the ROS production by inflammatory monocytes *ex vivo*. Our data indicated that *Cblb*^*+/+*^ cells had lesser ROS expression compared with *Cblb*^*-/-*^ (**Fig7C**). Thus, CBLB negatively regulates the ROS production in inflammatory monocytes, which was reminiscent of ROS production by BMMs *in vitro* (**Fig. 3G & H**). Collectively, our data show that inflammatory monocyte activation, but not ROS production, was inhibited in the absence of CBLB.

**Figure 7:**
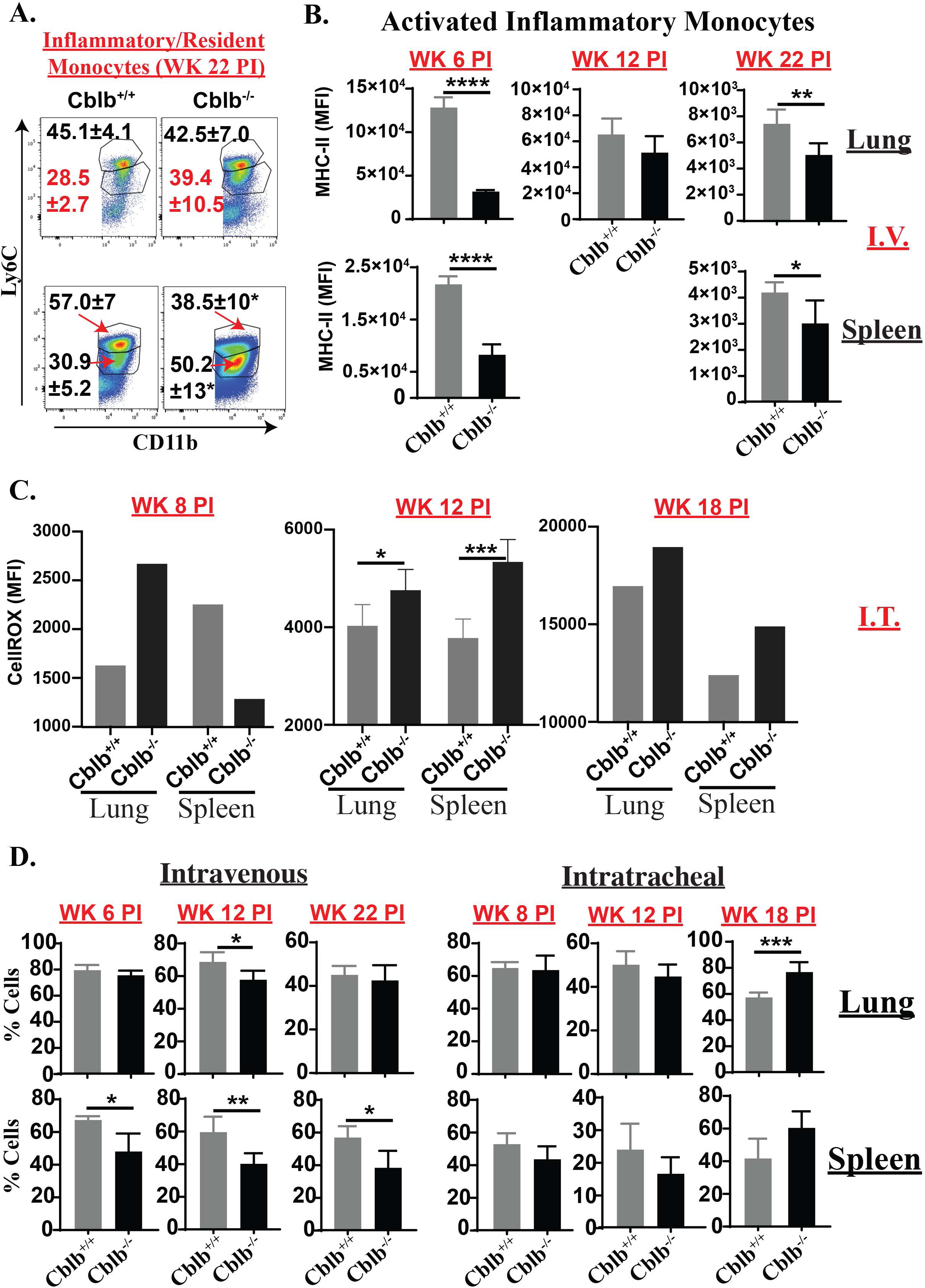
Abnormal Monocytes responses under *Cblb*-deficiency. *Cblb*^*+/+*^and *Cblb*^*-/-*^ mice were infected as described in Fig. 2. At indicated week PI, tissues were harvested, single-cell suspensions were stained for Inflammatory/Resident Monocytes (CD90^-^, Ly6G^-^, NK1.1^-^, CD11b^+^, Siglec-F^-^, Ly6C^hi/+^, respectively) using fluorochrome-conjugated antibodies, and analyzed by flow cytometry. (**A**). Plots show the percent of Inflammatory or Resident Monocytes. **B.** MFI of MHC-II expression. (**C**). MFI of CellROX (pooled or individual mouse samples). (**D**) Frequencies of inflammatory monocytes. Values are mean ± SD. N=3-6 mice/group. IV infection data is representative of 2-3 independent experiments. *p≤0.05, **p≤0.01, ***p≤0.001, and ***p≤0.001.

### Role of CBLB in neutrophils during an NTM infection

The neutrophil responses during mycobacterial diseases is a double-edged sword, and thcontribution to immunity is debated (Lowe et al., 2012; Lyadova, 2017). Here, we assessed neutrophil responses during NTM infection. First, we examined if the phagocytic ability of neutrophils was affected by CBLB. Bone-marrow-derived neutrophils (BMN) were incubated with 5MOI of dsRED^+^MAV104, and analyzed by flow cytometry. At 4hr and 12hr PI, ∼50% and ∼90%, respectively, of both *Cblb*^*+/+*^ and *Cblb*^*-/-*^ BMN were dsRED^+^ (**Fig. 8A**), suggesting a minimal role of CBLB for phagocytosis. However, there was a significant increase in activation status (MFI of CD44^hi^) of *Cblb*^*-/-*^ BMN compared with *Cblb*^*+/+*^ cells (**Fig. 8B**). Next, we evaluated neutrophil numbers *in vivo*. **Fig. 8C** shows the gating of frequencies of neutrophils in flow plots in the lung and spleen at Wk 22 PI, and found no differences between the groups. Following *i.v.* infection, neutrophil numbers were significantly reduced at an early time point in the spleen (Wk 6) in *Cblb*^*-/-*^ mice compared with *Cblb*^*+/+*^ (**Fig. 8E; bottom left panel**). However, in other time points and in the lungs, CBLB played a minimal role for neutrophil numbers (**Fig. 8E; left panels**). Similarly, CBLB was dispensable in regulating the neutrophil numbers following *i.t.* infection, except that we found a significant difference in the lung at Wk 18 PI. Next, we asked if neutrophil activation, which may be involved in pathology, was modulated by CBLB. Interestingly, on the contrary to *in vitro* data, we found significantly lower MFI of CD44 on neutrophils of *Cblb*^*-/-*^ compared with *Cblb*^*+/+*^ mice (**Fig. 8D**). Similarly, we found defect in activation of neutrophils in the absence of *Cblb* in lung and spleen, following i.v. and i.t routes of infection, respectively (**Supp. Fig. 3A & B**). Collectively, CBLB may be redundant for neutrophil numbers *in vivo*, but may help maintain the activation status during an NTM infection.

**Figure 8:**
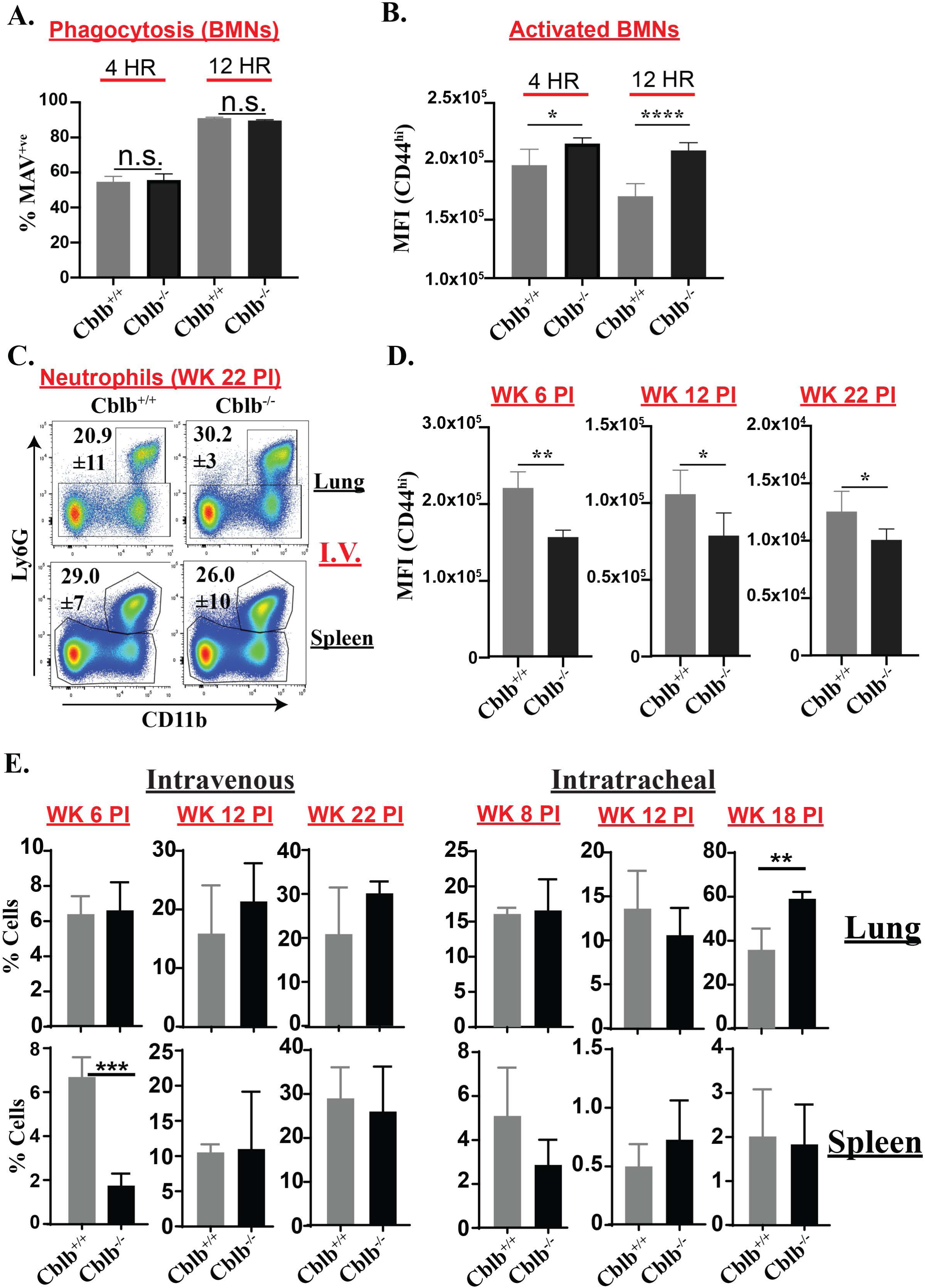
Neutrophil responses under *Cblb*-deficiency. **A & B**. Bone marrow-derived neutrophils (BMN) were infected with 5MOI of dsRED^+^MAV104. At indicated time points, rate of phagocytosis (% dsRED^+ve^; **A**) and activation (MFI of CD44; **B**) of neutrophils were analyzed by flow cytometry. (**C-E**) Mice were infected as described in Fig. 2. At indicated week PI, tissues were harvested, single-cell suspensions were stained for Neutrophils (CD90^-^, Ly6G^+^, CD11b^+^, NK1.1^-^) using fluorochrome-conjugated antibodies, and analyzed by flow cytometry. Dot plots (**C**) and Bar diagrams (**E**) show the percent of cells. **D.** Bar diagrams show the MFI of CD44 expression. IV infection data is representative of 2-3 independent experiments. Values are mean ± SD. N=3-6 mice/group. *p≤0.05, **p≤0.01, ***p≤0.001 and ****p≤0.0001.

### Dynamics of eosinophil numbers in the absence of *Cblb* during an NTM infection

Next, we asked if CBLB promotes an inadvertent response that may have affected bacterial growth and immunity (Kirman et al., 2000; Pfeffer et al., 2017). Following *i.v.* infection, we observed a significant increase in eosinophil numbers in *Cblb*^*-/-*^ mice compared with *Cblb*^*+/+*^ group (**Fig. 9A**). Following *i.t.* infection, eosinophil responses were significantly higher in *Cblb*^*-/-*^ mice compared with *Cblb*^*+/+*^ group at Wk 8PI in the lung, but not during later time points of infection (**Fig. 9B**). In the spleens, responses were more dynamic, but no significant differences between *Cblb*^*-/-*^ and *Cblb*^*+/+*^ mice were observed in the later stages of infection (**Fig. 9B**). Overall, our data suggested that CBLB plays a minimal or negative role in the regulation of eosinophil numbers following NTM infection.

**Figure 9:**
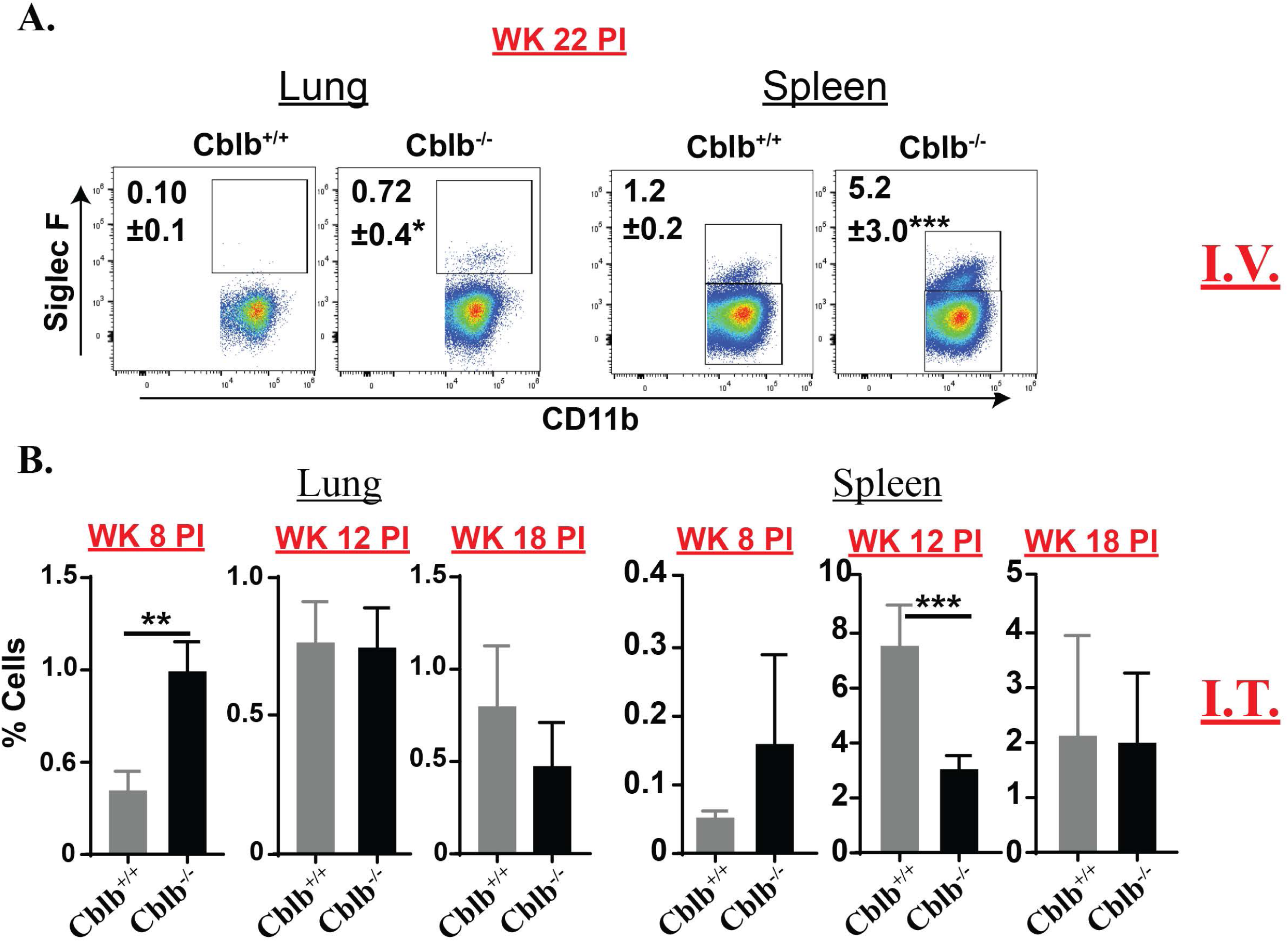
Eosinophil numbers in the absence of *Cblb*. Mice were infected as described in Fig. 2. At indicated week PI, tissues were harvested, single-cell suspensions were stained for Eosinophils (CD90^-^, Ly6G^-^, NK1.1^-^, CD11c^-^, CD11b^+^, Siglec-F^+^) using fluorochrome-conjugated antibodies, and analyzed by flow cytometry. **A.** Plots show the percent of cells following IV infection. **B.** Bar diagrams show the frequencies of eosinophils in the lung and spleens of mice infected by I.T. route. IV infection data is representative of 2 independent experiments. Values are mean ± SD. N=3-6 mice/group. *p≤0.05, **p≤0.01, and ***p≤0.001.

### CBLB alters the Plasmacytoid DC numbers during an NTM infection

Although the *in vivo* source of type I IFNs has yet to be clearly defined during NTM infection, myeloid cells, including plasmacytoid DC (pDC) cells, are known to be significant producers. Type I IFNs modulate disease pathogenesis and host responses during mycobacterial infection, both negatively and positively (Donovan et al., 2017; Lu et al., 2017; Moreira-Teixeira et al., 2018; Parlato et al., 2018). Here, we enumerated the number of plasmacytoid DC during an NTM infection. Following *i.v.* infection, plasmacytoid DC were significantly lower in the absence of *Cblb* at both Wks 6 and 12 PI (**Fig. 10B**), but not at a later time point (Wk 22PI; **Fig. 10A & B**). However, following *i.t.* infection, plasmacytoid DC were not affected, except at Wk12 PI, where we found a significant reduction in *Cblb*^*-/-*^ compared with *Cblb*^*+/+*^ mice (**Fig. 10C**). Thus, our data suggest an early depletion or inhibition of pDC in the absence of *Cblb* during systemic infection at earlier time points.

**Figure 10:**
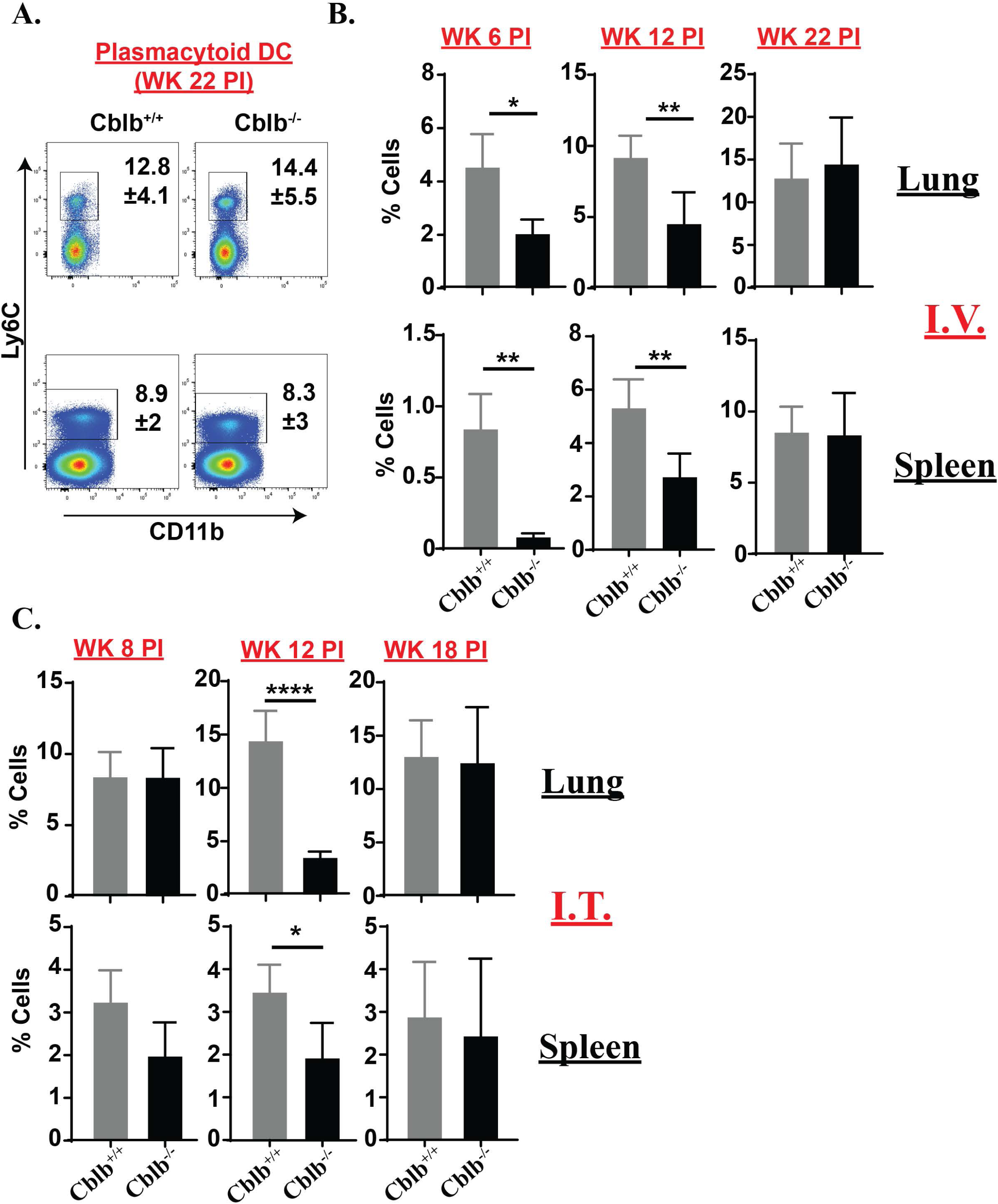
Plasmacytoid DC numbers under *Cblb*-deficiency. *Cblb*^*+/+*^and *Cblb*^*-/-*^ mice were infected as described in Fig. 2. At indicated week PI, tissues were harvested, single-cell suspensions were stained for Plasmacytoid DC (CD90^-^, Ly6G^-^, NK1.1^-^, CD11c^-^, CD11b^-^, Ly6C^+^) using fluorochrome-conjugated antibodies, and analyzed by flow cytometry. **A.** Dot plots show the percent of cells. **B & C.** Bar diagrams show the frequencies of plasmacytoid DC in the lung and spleens of mice infected by I.V. and I.T. routes, respectively. IV infection data is representative of 2-3 independent experiments. Values are mean ± SD. N=3-6 mice/group. *p≤0.05, **p≤0.01, and ****p≤0.0001.

### CBLB is required for conventional dendritic cell responses during an NTM infection

Next, we evaluated the role of CBLB for conventional DC (cDC) responses, which are necessary for productive adaptive immunity, and modulation of inflammation during mycobacterial infections (Mihret, 2012; Lai et al., 2018b; Ribechini et al., 2019). We measured the numbers of cDC in the spleen (both CD8^+^/CD11b^-^ and CD8^-^/CD11b^+^), and found that *Cblb*^*-/-*^ mice had significantly blunted cDC numbers at all the time points compared with *Cblb*^*+/+*^ in (**Fig. 11A & B**). Next, we asked if level of MHC-II, a classical marker for cDC activation, was affected by CBLB. Similar to the numbers, activation status (MFI of MHC-II) on CD8^+^ cDC (cross-presenting cells), but not on CD8^-^ cDC, were significantly reduced in the absence of *Cblb* at earlier time points of infection (**Fig. 11C**; data not shown). Thus, CBLB helps for robust cDC responses during an NTM infection.

**Figure 11:**
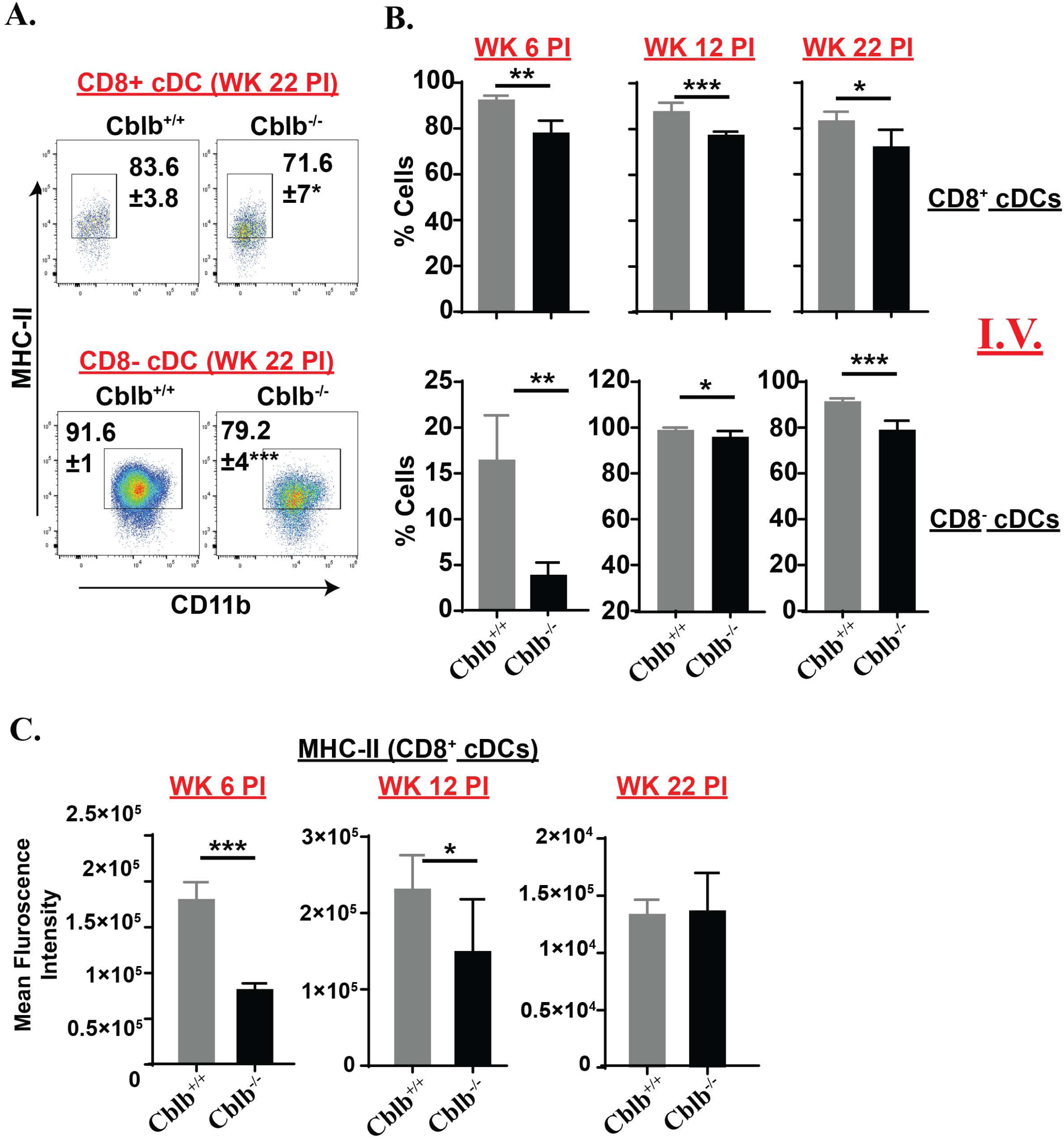
Conventional DC responses in *Cblb*-deficient mice. *Cblb*^*+/+*^and *Cblb*^*-/-*^ mice were i.v. infected as described in Fig. 2. At indicated week PI, spleens were harvested, single-cell suspensions were stained for cDC (CD90^-^, Ly6G^-^, NK1.1^-^, MHC-II^+^, CD11b^+^/^-^) using fluorochrome-conjugated antibodies, and analyzed by flow cytometry. **A.** Dot plots show the percent of cells. **B & C.** Bar diagrams show the frequencies of plasmacytoid DC. The data is representative of 2-3 independent experiments. Values are mean ± SD. N=3-6 mice/group. *p≤0.05, **p≤0.01, and ***p≤0.001.

### Role of CBLB in granulomatous inflammation during an NTM infection

Next, we wanted to determine the effect of CBLB for NTM pathogenesis by histopathology. Although the beneficial role of granuloma formation for the host defense is debated (Ramakrishnan, 2012; Silva Miranda et al., 2012; Pagan and Ramakrishnan, 2014), the dogma, nevertheless, supports for prevention of dissemination (Saunders and Cooper, 2000; Ndlovu and Marakalala, 2016). We dissected the role of *Cblb* for granulomatous inflammation, orchestrated by innate immune cells under T-cell deficiency. We harvested tissues following an NTM infection at Wks 3, 6, 9, and 13, and subjected for histopathological readings. The inflammation, characterized by small discrete granulomatous foci, was prominent in *Cblb*^*+/+*^ mice at 3-wk PI in the spleen & liver (**Fig. 12**). Similarly, multiple small indiscrete granulomatous foci were found in the lungs. In contrast, granulomatous foci were not apparent in the *Cblb*^*-/-*^ mice in any of the tissues at 3-wk PI and a few at 9-wk PI. However, small discrete granulomatous inflammation foci were observed at 13-wk PI in all tissues in *Cblb*^*-/-*^ mice (**Fig. 12**). These readings suggest a delay in the formation of granulomas in the absence of *Cblb*. Many increasing numbers of acid-fast bacilli, within the cytoplasm of macrophages, were noticed in *Cblb*^*-/-*^ mice compared with *Cblb*^*+/+*^ starting from 3-wk PI (data not shown) that was reminiscent of CFU readings (**Fig. 2**). Collectively, our data suggest that CBLB facilitated the early formation of granulomas and thwarting of dissemination during an NTM infection under T-cell deficiency.

**Figure 12:**
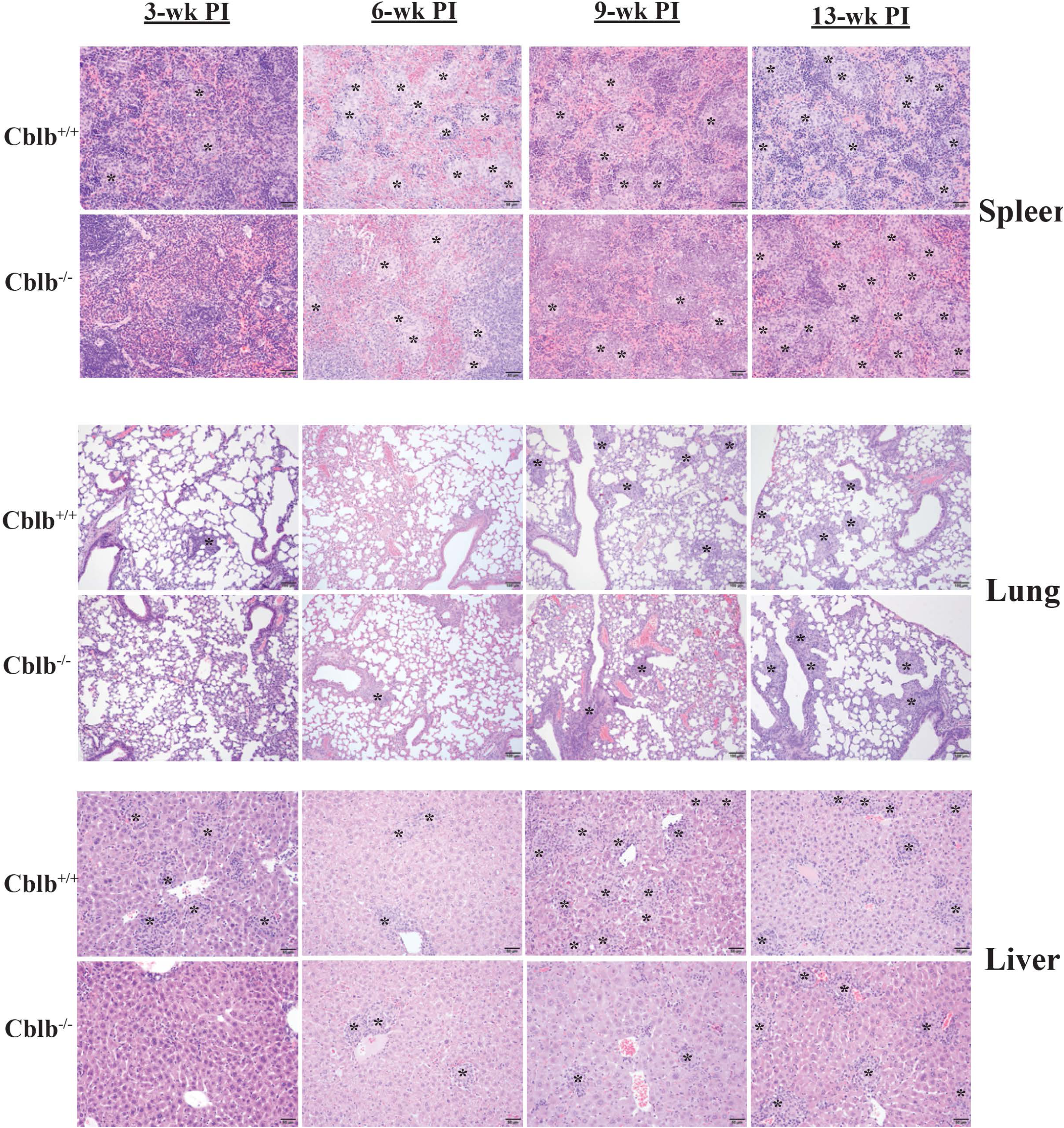
Histopathology of infected tissues. *Cblb*^*+/+*^and *Cblb*^*-/-*^ mice were infected as described in Fig. 2 by *i.v.* route. At indicated week PI, tissues were harvested in 10% buffered Neutral Formalin. Tissues were paraffin embedded, sectioned and mounted on the slides to stain with Hematoxylin and Eosin (H&E). The multiple samples were read at multiple foci for histopathological readouts, and representative photomicrographs of granulomatous inflammatory foci (asterisks) are shown here. Bar=50μm (Liver and Spleen), and 100μm (Lung). N=5-6 mice/group. Data is from 2-3 independent experiments.

## Discussion

Host regulators of innate immune responses during NTM infections are not well studied, including under severe T-cell deficiency. In this study, we systematically evaluated the role of CBLB for innate immunity and control of bacterial dissemination during an NTM infection in mice. We found that *Cblb*-deficiency enhanced bacterial growth and dissemination *in vivo*. We found several altered or defective innate immune cells under CBLB deficiency, including reduced NK cell numbers, reduced activation/numbers of inflammatory monocytes, conventional Dendritic Cells, and neutrophils. Notably, our studies showed that CBLB facilitated the early induction of granulomatous inflammation. Additionally, we showed the induction of CBLB following NTM infection, and *Cblb*-deficient macrophages were competent to control the growth of bacteria.

Mouse models of NTM infections, especially using C57BL/6 and *M. avium*, suggest that bacterial pathogenesis can be studied [(Chan ED et al., Mycobact Dis 2016, 6:3); (Verma et al., 2019)]. Here we define CBLB as an important innate host regulators of NTM infections. Our study showed that CBLB is necessary for inhibiting NTM growth and dissemination under T-cell deficiency (**Fig. 2**). To delineate the cellular mechanisms, we sought to determine if CBLB could be induced in macrophages, primary target cells of mycobacteria, following NTM infection. We discovered that CBLB was significantly induced in primary macrophages, including splenocytes. Next, we delineated the role of CBLB for NTM growth in macrophages *in vitro* using BMM, and found that *Cblb*^*-/-*^ cells were equally competent or superior in inhibiting NTM growth compared with *Cblb*^*+/+*^ (**Fig. 3**), which may be due to enhanced ROS production (Shastri et al., 2018). Our *in vitro* immunity (CFU) data was in line with other studies using fungi (Wirnsberger et al., 2016; Xiao et al., 2016), but was in contrast to recently published report using M. *tuberculosis* (Penn et al., 2018). The latter study suggested that CBLB was required to sequester LpqN, an Mtb virulence factor, for enhanced bacterial killing by macrophages. Interestingly, in this study, lpqN mutant Mtb was still able to grow higher in *Cblb*-deficient macrophages compared with *Cblb*-sufficient cells. Although utative LpqN family protein is present in M. *avium* (NCBI blast, sequence ID: EUA40567.1 and Broad institute MAV4561 protein) species, the lifestyle/pathogenesis of these bacteria may be different in macrophages. Further studies are warranted to decipher the *in vivo* role of CBLB in macrophages during mycobacterial diseases.

NK cells are critical regulators of pathogenesis during mycobacterial, including NTM infections (Dhiman et al., 2012; Allen et al., 2015; Lai et al., 2018a). Our study showed a significant loss of NK cells in *Cblb*-deficient mice, especially of CD11b^+^ subset. To validate our observations on the role of CBLB in NK cells *in vivo*, we investigated if NTM infection would affect the expression of CBLB and its function using *in vitro*. NTM infection did significantly enhance CBLB levels in NK cells, and as predicted (Liu et al., 2014), *Cblb*^-/-^ NK cells produced higher IFNγ compared with *Cblb*^+/+^ (**Fig. 5**). Hence, we postulate that impaired NK cell numbers in *Cblb-*deficient mice may be one of the causes of higher bacterial burden and dissemination. Our results are in line with the potential beneficial role of NK cells to control NTM infection (Lai et al., 2018a), and CD11b^+^ NK cells may be of superior in functions (Lu et al., 2014; Allen et al., 2015; Venkatasubramanian et al., 2017; Garand et al., 2018; Lai et al., 2018a). Although the role of *Cblb*-deficient NK cells during the early phases of infection and their loss during NTM infection is not known, we think a higher antigen level is the probable cause for their depletion, akin to NKT or T cells (Fuller and Zajac, 2003; Kee et al., 2012). Further studies are required to determine CBLB-dependent NK cells for control of NTM infection.

There is an immense interest in the role of newly recruited monocytes, including inflammatory monocytes, in the pathogenesis of mycobacterial diseases (Serbina et al., 2008; Marakalala et al., 2018; Sampath et al., 2018; Davis et al., 2019). Monocytes/macrophages, recently recruited, expressing CCR2 seem to be highly permissive to *M. tuberculosis* due to their inability to get activation signals that are masked by phenolic glycolipids decoration on bacteria (Cambier et al., 2017; Garcia-Vilanova et al., 2019). However, several studies have shown the enhanced susceptibility under CCR2-deficiency, which may also depend on the virulence of mycobacteria (Peters et al., 2001; Dunlap et al., 2018; Garcia-Vilanova et al., 2019). Thus, newly recruited monocytes are critical regulators of pathogenesis and outcome of mycobacterial diseases, and can be targeted for therapeutic interventions (Norris and Ernst, 2018). Additionally, monocytes/macrophages can regulate defense against mycobacteria by producing reactive oxygen/nitrogen species (Lamichhane, 2011), and priming productive T-cell responses (Scott and Flynn, 2002; Peters et al., 2004; Samstein et al., 2013). In our study, we found a poor activation (low MHC-II), but higher production of ROS, in *Cblb*-deficient inflammatory monocytes (**Fig. 7**). However, numbers of inflammatory monocytes were relatively intact, except in the spleens of IV infected groups, where we found enhanced numbers of non-inflammatory monocytes. These non-inflammatory monocytes, in a recent study, are classified as Myeloid-Derived Suppressor Cells, subvert T-cell responses and cause impaired DC response (Abdissa et al., 2018). We reasoned if CBLB regulates the expression of *CCL2*, a chemokine involved in the recruitment of monocytes. We found either normal or less *CCL2* expression in infected *Cblb*^-/-^ cells compared with *Cblb*^*+/+*^, *in vitro* (**Supp. Fig. 4**), suggesting a minimal role of CCL2 for the abnormal phenotype. Nevertheless, *in vivo* dynamics of CCL2 expression and its role in the activation of monocytes during NTM infection needs further investigation (Bose and Cho, 2013; Domingo-Gonzalez et al., 2016). The low MHC-II expression levels on *Cblb*-deficient inflammatory monocytes may be due to their less differentiation status or commitment into macrophage/dendritic cell (**Fig. 6B & 11**) lineage under altered micromilieu (Rivollier et al., 2012; Srivastava et al., 2014; Lastrucci et al., 2015; Sampath et al., 2018). Nevertheless, we found an enhanced ROS production under *Cblb*-deficiency *in vivo* that recapitulated our *in vitro* BMM studies (**Fig. 3**), and may, perhaps, be involved in dissemination in the lack of T-cell help or enhanced cell death (Roca and Ramakrishnan, 2013; Divangahi et al., 2018).

Neutrophils play a dichotomous role during mycobacterial infections, and their role for contributing to immunity or immunopathology is confounded (Lowe et al., 2012; Kroon et al., 2018). Early recruitment and status of neutrophils seem to dictate their role in the control the mycobacterial infection, and many studies show the importance of neutrophil function in CGD patients in defense against active tuberculosis (Martineau et al., 2007; Kulkarni et al., 2016; Mishra et al., 2017; Wolach et al., 2017; Lowe et al., 2018; Gideon et al., 2019). However, exuberant neutrophil responses are associated with pathology (Lyadova, 2017). Our data suggested that phagocytosis of bacteria by neutrophils was not affected by *Cblb*-deficiency. However, CBLB seems to inhibit the activation of BMN (MFI of CD44) *in vitro*. However, our *in vivo* studies suggested otherwise, in that, the activation status of neutrophils was significantly less in *Cblb*-deficient mice, but numbers were not affected. Similarly, we did not see significant differences in the numbers of neutrophils among the groups by histopathology, despite increased numbers of bacteria in the macrophages of *Cblb*-deficient groups. Although the role of CBLB for neutrophil functions is understudied, a recent study suggested a minimal effect of CBLB on neutrophil responses during candida infection (Xiao et al., 2016). Further functional studies are required to delineate CBLB role in neutrophils during the early phases of NTM infection. (Appelberg et al., 1995; Lake et al., 2016).

Dendritic cells are of diverse types, but we classified them into two major groups in our study. Plasmacytoid DC-immune modulators with little direct antigen presentation; and conventional DC-which have a direct impact on adaptive immunity with antigen-presentation or activation (Mihret, 2012; Parlato et al., 2018). In our study, we found a significant reduction in plasmacytoid and conventional DC responses under *Cblb*-deficiency. We postulate that defective DC cells in *Cblb*-deficient mice may have caused the dysfunction of other innate immune cells. The defective DC responses may be due to lack of Th1 responses (Frasca et al., 2008) and, in the absence of such, with impaired clearance of bacteria, Th2 polarization may occur (Traynor et al., 2000; Mendez-Samperio, 2010; Pfeffer et al., 2017). In line with this, we found an exuberant eosinophil numbers in the absence of *Cblb*. Eosinophil numbers, an indicator of Th2 responses, were higher in spleens following i.v. infection, especially in later phases. Studies in mycobacterial diseases suggested a negative role of eosinophils (Pfeffer et al., 2017; Moideen et al., 2018), and their depletion fostered the immunity, but not prevention of dissemination (Kirman et al., 2000). We could not explain the differences in the eosinophil numbers following *i.v./i.t*. routes of infection, but we believe that it may be due to higher CFUs following i.v. infection.

In this study, we have shown that CBLB was necessary for controlling bacterial growth and dissemination during an NTM infection under compromised T-cell immunity; the deficiency led to poor or altered status of many innate immune cell subsets, and the lack of early granulomatous inflammation. Future studies are warranted to uncover the role of CBLB in each innate immune cell subsets during NTM infection. Collectively, our studies demonstrate that CBLB can be a target for therapeutic and preventive measures in controlling NTM infections.

## Acknowledgments

We sincerely thank Dr. Howard Steinberg, Dept. of Pathobiological Sciences at the University of Wisconsin-Madison for some of the histopathological studies, and Dr. Gee W. Lau, Dept. of Pathobiology at the UIUC for CBLB protein analysis experiments. We thank animal care facility at the University of Illinois at Urbana-Champaign. We also thank many investigators who have contributed significantly in the field of NTM or mycobacterial diseases that were missed in citations.

## Author Contributions

SM and JS designed the experiments, executed, and analyzed the data. HFMA designed, executed, and analyzed some of the *in vitro* & *in vivo* experiments. TCK executed and analyzed some *in vitro* experiments. WC designed, executed, and analyzed western blot experiments. AMT provided the Mycobacterium strains and helped in designing some of the experiments. MDV analyzed the histopathological data and edited the MS. SGN conceived the project, designed the experiments, executed and analyzed the data, and wrote the manuscript.

## Ethics Statement

This work was carried in accordance with the protocol approved by IACUC committee at the University of Illinois at Urbana-Champaign.

## References

Abdissa, K., Nerlich, A., Beineke, A., Ruangkiattikul, N., Pawar, V., Heise, U., et al. (2018). Presence of Infected Gr-1(int)CD11b(hi)CD11c(int) Monocytic Myeloid Derived Suppressor Cells Subverts T Cell Response and Is Associated With Impaired Dendritic Cell Function in Mycobacterium avium-Infected Mice. Front Immunol 9, 2317. doi: 10.3389/fimmu.2018.02317.

Adjemian, J., Olivier, K.N., Seitz, A.E., Holland, S.M., and Prevots, D.R. (2012). Prevalence of nontuberculous mycobacterial lung disease in U.S. Medicare beneficiaries. Am J Respir Crit Care Med 185(8), 881–886. doi: 10.1164/rccm.201111-2016OC.

Allen, M., Bailey, C., Cahatol, I., Dodge, L., Yim, J., Kassissa, C., et al. (2015). Mechanisms of Control of Mycobacterium tuberculosis by NK Cells: Role of Glutathione. Front Immunol 6, 508. doi: 10.3389/fimmu.2015.00508.

Appelberg, R., Castro, A.G., Gomes, S., Pedrosa, J., and Silva, M.T. (1995). Susceptibility of beige mice to Mycobacterium avium: role of neutrophils. Infect Immun 63(9), 3381–3387.

Arron, J.R., Vologodskaia, M., Wong, B.R., Naramura, M., Kim, N., Gu, H., et al. (2001). A positive regulatory role for Cbl family proteins in tumor necrosis factor-related activation-induced cytokine (trance) and CD40L-mediated Akt activation. J Biol Chem 276(32), 30011–30017. doi: 10.1074/jbc.M100414200.

Bachmaier, K., Krawczyk, C., Kozieradzki, I., Kong, Y.Y., Sasaki, T., Oliveira-dos-Santos, A., et al. (2000). Negative regulation of lymphocyte activation and autoimmunity by the molecular adaptor Cbl-b. Nature 403(6766), 211–216. doi: 10.1038/35003228.

Bachmaier, K., Toya, S., Gao, X., Triantafillou, T., Garrean, S., Park, G.Y., et al. (2007). E3 ubiquitin ligase Cblb regulates the acute inflammatory response underlying lung injury. Nat Med 13(8), 920–926. doi: 10.1038/nm1607.

Behar, S.M., Divangahi, M., and Remold, H.G. (2010). Evasion of innate immunity by Mycobacterium tuberculosis: is death an exit strategy? Nat Rev Microbiol 8(9), 668–674. doi: 10.1038/nrmicro2387.

Bose, S., and Cho, J. (2013). Role of chemokine CCL2 and its receptor CCR2 in neurodegenerative diseases. Arch Pharm Res 36(9), 1039–1050. doi: 10.1007/s12272-013-0161-z.

Cambier, C.J., O’Leary, S.M., O’Sullivan, M.P., Keane, J., and Ramakrishnan, L. (2017). Phenolic Glycolipid Facilitates Mycobacterial Escape from Microbicidal Tissue-Resident Macrophages. Immunity 47(3), 552–565 e554. doi: 10.1016/j.immuni.2017.08.003.

Chiang, Y.J., Kole, H.K., Brown, K., Naramura, M., Fukuhara, S., Hu, R.J., et al. (2000). Cbl-b regulates the CD28 dependence of T-cell activation. Nature 403(6766), 216–220. doi: 10.1038/35003235.

Chirino, L.M., Kumar, S., Okumura, M., Sterner, D.E., Mattern, M., Butt, T.R., et al. (2019). TAM receptors attenuate murine NK-cell responses via E3 ubiquitin ligase Cbl-b. Eur J Immunol. doi: 10.1002/eji.201948204.

Cohen, S.B., Gern, B.H., Delahaye, J.L., Adams, K.N., Plumlee, C.R., Winkler, J.K., et al. (2018). Alveolar Macrophages Provide an Early Mycobacterium tuberculosis Niche and Initiate Dissemination. Cell Host Microbe 24(3), 439–446 e434. doi: 10.1016/j.chom.2018.08.001.

Cong, J., and Wei, H. (2019). Natural Killer Cells in the Lungs. Front Immunol 10, 1416. doi: 10.3389/fimmu.2019.01416.

Davis, A.G., Rohlwink, U.K., Proust, A., Figaji, A.A., and Wilkinson, R.J. (2019). The pathogenesis of tuberculous meningitis. J Leukoc Biol 105(2), 267–280. doi: 10.1002/JLB.MR0318-102R.

DeWan, A.T., Egan, K.B., Hellenbrand, K., Sorrentino, K., Pizzoferrato, N., Walsh, K.M., et al. (2012). Whole-exome sequencing of a pedigree segregating asthma. BMC Med Genet 13, 95. doi: 10.1186/1471-2350-13-95.

Dhiman, R., Periasamy, S., Barnes, P.F., Jaiswal, A.G., Paidipally, P., Barnes, A.B., et al. (2012). NK1.1+ cells and IL-22 regulate vaccine-induced protective immunity against challenge with Mycobacterium tuberculosis. J Immunol 189(2), 897–905. doi: 10.4049/jimmunol.1102833.

Diel, R., Lipman, M., and Hoefsloot, W. (2018). High mortality in patients with Mycobacterium avium complex lung disease: a systematic review. BMC Infect Dis 18(1), 206. doi: 10.1186/s12879-018-3113-x.

Divangahi, M., Khan, N., and Kaufmann, E. (2018). Beyond Killing Mycobacterium tuberculosis: Disease Tolerance. Front Immunol 9, 2976. doi: 10.3389/fimmu.2018.02976.

Domingo-Gonzalez, R., Prince, O., Cooper, A., and Khader, S.A. (2016). Cytokines and Chemokines in Mycobacterium tuberculosis Infection. Microbiol Spectr 4(5). doi: 10.1128/microbiolspec.TBTB2-0018-2016.

Doniz-Padilla, L., Martinez-Jimenez, V., Nino-Moreno, P., Abud-Mendoza, C., Hernandez-Castro, B., Gonzalez-Amaro, R., et al. (2011). Expression and function of Cbl-b in T cells from patients with systemic lupus erythematosus, and detection of the 2126 A/G Cblb gene polymorphism in the Mexican mestizo population. Lupus 20(6), 628–635. doi: 10.1177/0961203310394896.

Donovan, M.L., Schultz, T.E., Duke, T.J., and Blumenthal, A. (2017). Type I Interferons in the Pathogenesis of Tuberculosis: Molecular Drivers and Immunological Consequences. Front Immunol 8, 1633. doi: 10.3389/fimmu.2017.01633.

Dorhoi, A., Yeremeev, V., Nouailles, G., Weiner, J., 3rd, Jorg, S., Heinemann, E., et al. (2014). Type I IFN signaling triggers immunopathology in tuberculosis-susceptible mice by modulating lung phagocyte dynamics. Eur J Immunol 44(8), 2380–2393. doi: 10.1002/eji.201344219.

Dunlap, M.D., Howard, N., Das, S., Scott, N., Ahmed, M., Prince, O., et al. (2018). A novel role for C-C motif chemokine receptor 2 during infection with hypervirulent Mycobacterium tuberculosis. Mucosal Immunol 11(6), 1727–1742. doi: 10.1038/s41385-018-0071-y.

Frasca, L., Nasso, M., Spensieri, F., Fedele, G., Palazzo, R., Malavasi, F., et al. (2008). IFN-gamma arms human dendritic cells to perform multiple effector functions. J Immunol 180(3), 1471–1481. doi: 10.4049/jimmunol.180.3.1471.

Fu, B., Tian, Z., and Wei, H. (2014). Subsets of human natural killer cells and their regulatory effects. Immunology 141(4), 483–489. doi: 10.1111/imm.12224.

Fuller, M.J., and Zajac, A.J. (2003). Ablation of CD8 and CD4 T cell responses by high viral loads. J Immunol 170(1), 477–486. doi: 10.4049/jimmunol.170.1.477.

Garand, M., Goodier, M., Owolabi, O., Donkor, S., Kampmann, B., and Sutherland, J.S. (2018). Functional and Phenotypic Changes of Natural Killer Cells in Whole Blood during Mycobacterium tuberculosis Infection and Disease. Front Immunol 9, 257. doi: 10.3389/fimmu.2018.00257.

Garcia-Vilanova, A., Chan, J., and Torrelles, J.B. (2019). Underestimated Manipulative Roles of Mycobacterium tuberculosis Cell Envelope Glycolipids During Infection. Front Immunol 10, 2909. doi: 10.3389/fimmu.2019.02909.

Gideon, H.P., Phuah, J., Junecko, B.A., and Mattila, J.T. (2019). Neutrophils express pro- and anti-inflammatory cytokines in granulomas from Mycobacterium tuberculosis-infected cynomolgus macaques. Mucosal Immunol 12(6), 1370–1381. doi: 10.1038/s41385-019-0195-8.

Grainger, J.R., Wohlfert, E.A., Fuss, I.J., Bouladoux, N., Askenase, M.H., Legrand, F., et al. (2013). Inflammatory monocytes regulate pathologic responses to commensals during acute gastrointestinal infection. Nat Med 19(6), 713–721. doi: 10.1038/nm.3189.

Henkle, E., and Winthrop, K.L. (2015). Nontuberculous mycobacteria infections in immunosuppressed hosts. Clin Chest Med 36(1), 91–99. doi: 10.1016/j.ccm.2014.11.002.

Heung, L.J., and Hohl, T.M. (2019). Inflammatory monocytes are detrimental to the host immune response during acute infection with Cryptococcus neoformans. PLoS Pathog 15(3), e1007627. doi: 10.1371/journal.ppat.1007627.

Hey, Y.Y., Quah, B., and O’Neill, H.C. (2017). Antigen presenting capacity of murine splenic myeloid cells. BMC Immunol 18(1), 4. doi: 10.1186/s12865-016-0186-4.

Horne, D., and Skerrett, S. (2019). Recent advances in nontuberculous mycobacterial lung infections. F1000Res 8. doi: 10.12688/f1000research.20096.1.

International Multiple Sclerosis Genetics, C., Hafler, D.A., Compston, A., Sawcer, S., Lander, E.S., Daly, M.J., et al. (2007). Risk alleles for multiple sclerosis identified by a genomewide study. N Engl J Med 357(9), 851–862. doi: 10.1056/NEJMoa073493.

Jeon, M.S., Atfield, A., Venuprasad, K., Krawczyk, C., Sarao, R., Elly, C., et al. (2004). Essential role of the E3 ubiquitin ligase Cbl-b in T cell anergy induction. Immunity 21(2), 167–177. doi: 10.1016/j.immuni.2004.07.013.

Johnson, M.M., and Odell, J.A. (2014). Nontuberculous mycobacterial pulmonary infections. J Thorac Dis 6(3), 210–220. doi: 10.3978/j.issn.2072-1439.2013.12.24.

Karwacz, K., Bricogne, C., MacDonald, D., Arce, F., Bennett, C.L., Collins, M., et al. (2011). PD-L1 co-stimulation contributes to ligand-induced T cell receptor down-modulation on CD8+ T cells. EMBO Mol Med 3(10), 581–592. doi: 10.1002/emmm.201100165.

Kee, S.J., Kwon, Y.S., Park, Y.W., Cho, Y.N., Lee, S.J., Kim, T.J., et al. (2012). Dysfunction of natural killer T cells in patients with active Mycobacterium tuberculosis infection. Infect Immun 80(6), 2100–2108. doi: 10.1128/IAI.06018-11.

Kirman, J., Zakaria, Z., McCoy, K., Delahunt, B., and Le Gros, G. (2000). Role of eosinophils in the pathogenesis of Mycobacterium bovis BCG infection in gamma interferon receptor-deficient mice. Infect Immun 68(5), 2976–2978. doi: 10.1128/iai.68.5.2976-2978.2000.

Koning, J.J., and Mebius, R.E. (2012). Interdependence of stromal and immune cells for lymph node function. Trends Immunol 33(6), 264–270. doi: 10.1016/j.it.2011.10.006.

Kosoy, R., Yokoi, N., Seino, S., and Concannon, P. (2004). Polymorphic variation in the CBLB gene in human type 1 diabetes. Genes Immun 5(3), 232–235. doi: 10.1038/sj.gene.6364057.

Kroon, E.E., Coussens, A.K., Kinnear, C., Orlova, M., Moller, M., Seeger, A., et al. (2018). Neutrophils: Innate Effectors of TB Resistance? Front Immunol 9, 2637. doi: 10.3389/fimmu.2018.02637.

Kulkarni, M., Desai, M., Gupta, M., Dalvi, A., Taur, P., Terrance, A., et al. (2016). Clinical, Immunological, and Molecular Findings of Patients with p47(phox) Defect Chronic Granulomatous Disease (CGD) in Indian Families. J Clin Immunol 36(8), 774–784. doi: 10.1007/s10875-016-0333-y.

Lai, H.C., Chang, C.J., Lin, C.S., Wu, T.R., Hsu, Y.J., Wu, T.S., et al. (2018a). NK Cell-Derived IFN-gamma Protects against Nontuberculous Mycobacterial Lung Infection. J Immunol 201(5), 1478–1490. doi: 10.4049/jimmunol.1800123.

Lai, R., Jeyanathan, M., Afkhami, S., Zganiacz, A., Hammill, J.A., Yao, Y., et al. (2018b). CD11b(+) Dendritic Cell-Mediated Anti-Mycobacterium tuberculosis Th1 Activation Is Counterregulated by CD103(+) Dendritic Cells via IL-10. J Immunol 200(5), 1746–1760. doi: 10.4049/jimmunol.1701109.

Lake, M.A., Ambrose, L.R., Lipman, M.C., and Lowe, D.M. (2016). ’“Why me, why now?” Using clinical immunology and epidemiology to explain who gets nontuberculous mycobacterial infection. BMC Med 14, 54. doi: 10.1186/s12916-016-0606-6.

Lamichhane, G. (2011). Mycobacterium tuberculosis response to stress from reactive oxygen and nitrogen species. Front Microbiol 2, 176. doi: 10.3389/fmicb.2011.00176.

Lastrucci, C., Benard, A., Balboa, L., Pingris, K., Souriant, S., Poincloux, R., et al. (2015). Tuberculosis is associated with expansion of a motile, permissive and immunomodulatory CD16(+) monocyte population via the IL-10/STAT3 axis. Cell Res 25(12), 1333–1351. doi: 10.1038/cr.2015.123.

Li, D., Gal, I., Vermes, C., Alegre, M.L., Chong, A.S., Chen, L., et al. (2004). Cutting edge: Cbl-b: one of the key molecules tuning CD28- and CTLA-4-mediated T cell costimulation. J Immunol 173(12), 7135–7139. doi: 10.4049/jimmunol.173.12.7135.

Li, P., Liu, H.L., Zhang, Z.Q., Lv, X.D., Chang, Y.X., Wang, H.J., et al. (2018). Single nucleotide polymorphisms of casitas B-lineage lymphoma proto-oncogene-b predict outcomes of patients with advanced non-small cell lung cancer after first-line platinum based doublet chemotherapy. J Thorac Dis 10(3), 1635–1647. doi: 10.21037/jtd.2018.02.31.

Liu, L., Wei, Y., and Wei, X. (2017). The Immune Function of Ly6Chi Inflammatory Monocytes During Infection and Inflammation. Curr Mol Med 17(1), 4–12. doi: 10.2174/1566524017666170220102732.

Liu, Q., Zhou, H., Langdon, W.Y., and Zhang, J. (2014). E3 ubiquitin ligase Cbl-b in innate and adaptive immunity. Cell Cycle 13(12), 1875–1884. doi: 10.4161/cc.29213.

Loeser, S., Loser, K., Bijker, M.S., Rangachari, M., van der Burg, S.H., Wada, T., et al. (2007). Spontaneous tumor rejection by cbl-b-deficient CD8+ T cells. J Exp Med 204(4), 879–891. doi: 10.1084/jem.20061699.

Loeser, S., and Penninger, J.M. (2007). Regulation of peripheral T cell tolerance by the E3 ubiquitin ligase Cbl-b. Semin Immunol 19(3), 206–214. doi: 10.1016/j.smim.2007.02.004.

Lowe, D.M., Demaret, J., Bangani, N., Nakiwala, J.K., Goliath, R., Wilkinson, K.A., et al. (2018). Differential Effect of Viable Versus Necrotic Neutrophils on Mycobacterium tuberculosis Growth and Cytokine Induction in Whole Blood. Front Immunol 9, 903. doi: 10.3389/fimmu.2018.00903.

Lowe, D.M., Redford, P.S., Wilkinson, R.J., O’Garra, A., and Martineau, A.R. (2012). Neutrophils in tuberculosis: friend or foe? Trends Immunol 33(1), 14–25. doi: 10.1016/j.it.2011.10.003.

Lu, C.C., Wu, T.S., Hsu, Y.J., Chang, C.J., Lin, C.S., Chia, J.H., et al. (2014). NK cells kill mycobacteria directly by releasing perforin and granulysin. J Leukoc Biol 96(6), 1119–1129. doi: 10.1189/jlb.4A0713-363RR.

Lu, Y.B., Xiao, D.Q., Liang, K.D., Zhang, J.A., Wang, W.D., Yu, S.Y., et al. (2017). Profiling dendritic cell subsets in the patients with active pulmonary tuberculosis. Mol Immunol 91, 86–96. doi: 10.1016/j.molimm.2017.08.007.

Lutz-Nicoladoni, C., Wolf, D., and Sopper, S. (2015). Modulation of Immune Cell Functions by the E3 Ligase Cbl-b. Front Oncol 5, 58. doi: 10.3389/fonc.2015.00058.

Lyadova, I.V. (2017). Neutrophils in Tuberculosis: Heterogeneity Shapes the Way? Mediators Inflamm 2017, 8619307. doi: 10.1155/2017/8619307.

Marakalala, M.J., Martinez, F.O., Pluddemann, A., and Gordon, S. (2018). Macrophage Heterogeneity in the Immunopathogenesis of Tuberculosis. Front Microbiol 9, 1028. doi: 10.3389/fmicb.2018.01028.

Martineau, A.R., Newton, S.M., Wilkinson, K.A., Kampmann, B., Hall, B.M., Nawroly, N., et al. (2007). Neutrophil-mediated innate immune resistance to mycobacteria. J Clin Invest 117(7), 1988–1994. doi: 10.1172/JCI31097.

Mendez-Samperio, P. (2010). Role of interleukin-12 family cytokines in the cellular response to mycobacterial disease. Int J Infect Dis 14(5), e366–371. doi: 10.1016/j.ijid.2009.06.022.

Mihret, A. (2012). The role of dendritic cells in Mycobacterium tuberculosis infection. Virulence 3(7), 654–659. doi: 10.4161/viru.22586.

Misharin, A.V., Morales-Nebreda, L., Mutlu, G.M., Budinger, G.R., and Perlman, H. (2013). Flow cytometric analysis of macrophages and dendritic cell subsets in the mouse lung. Am J Respir Cell Mol Biol 49(4), 503–510. doi: 10.1165/rcmb.2013-0086MA.

Mishra, B.B., Lovewell, R.R., Olive, A.J., Zhang, G., Wang, W., Eugenin, E., et al. (2017). Nitric oxide prevents a pathogen-permissive granulocytic inflammation during tuberculosis. Nat Microbiol 2, 17072. doi: 10.1038/nmicrobiol.2017.72.

Moideen, K., Kumar, N.P., Nair, D., Banurekha, V.V., Bethunaickan, R., and Babu, S. (2018). Heightened Systemic Levels of Neutrophil and Eosinophil Granular Proteins in Pulmonary Tuberculosis and Reversal following Treatment. Infect Immun 86(6). doi: 10.1128/IAI.00008-18.

Moreira-Teixeira, L., Mayer-Barber, K., Sher, A., and O’Garra, A. (2018). Type I interferons in tuberculosis: Foe and occasionally friend. J Exp Med 215(5), 1273–1285. doi: 10.1084/jem.20180325.

Nanjappa, S.G., Mudalagiriyappa, S., Fites, J.S., Suresh, M., and Klein, B.S. (2018). CBLB Constrains Inactivated Vaccine-Induced CD8(+) T Cell Responses and Immunity against Lethal Fungal Pneumonia. J Immunol 201(6), 1717–1726. doi: 10.4049/jimmunol.1701241.

Ndlovu, H., and Marakalala, M.J. (2016). Granulomas and Inflammation: Host-Directed Therapies for Tuberculosis. Front Immunol 7, 434. doi: 10.3389/fimmu.2016.00434.

Norris, B.A., and Ernst, J.D. (2018). Mononuclear cell dynamics in M. tuberculosis infection provide opportunities for therapeutic intervention. PLoS Pathog 14(10), e1007154. doi: 10.1371/journal.ppat.1007154.

Oh, S.Y., Park, J.U., Zheng, T., Kim, Y.K., Wu, F., Cho, S.H., et al. (2011). Cbl-b regulates airway mucosal tolerance to aeroallergen. Clin Exp Allergy 41(3), 434–442. doi: 10.1111/j.1365-2222.2010.03593.x.

Pagan, A.J., and Ramakrishnan, L. (2014). Immunity and Immunopathology in the Tuberculous Granuloma. Cold Spring Harb Perspect Med 5(9). doi: 10.1101/cshperspect.a018499.

Paolino, M., Choidas, A., Wallner, S., Pranjic, B., Uribesalgo, I., Loeser, S., et al. (2014). The E3 ligase Cbl-b and TAM receptors regulate cancer metastasis via natural killer cells. Nature 507(7493), 508–512. doi: 10.1038/nature12998.

Paolino, M., and Penninger, J.M. (2010). Cbl-b in T-cell activation. Semin Immunopathol 32(2), 137–148. doi: 10.1007/s00281-010-0197-9.

Paolino, M., Thien, C.B., Gruber, T., Hinterleitner, R., Baier, G., Langdon, W.Y., et al. (2011). Essential role of E3 ubiquitin ligase activity in Cbl-b-regulated T cell functions. J Immunol 186(4), 2138–2147. doi: 10.4049/jimmunol.1003390.

Parlato, S., Chiacchio, T., Salerno, D., Petrone, L., Castiello, L., Romagnoli, G., et al. (2018). Impaired IFN-alpha-mediated signal in dendritic cells differentiates active from latent tuberculosis. PLoS One 13(1), e0189477. doi: 10.1371/journal.pone.0189477.

Payne, F., Cooper, J.D., Walker, N.M., Lam, A.C., Smink, L.J., Nutland, S., et al. (2007). Interaction analysis of the CBLB and CTLA4 genes in type 1 diabetes. J Leukoc Biol 81(3), 581–583. doi: 10.1189/jlb.0906577.

Penn, B.H., Netter, Z., Johnson, J.R., Von Dollen, J., Jang, G.M., Johnson, T., et al. (2018). An Mtb-Human Protein-Protein Interaction Map Identifies a Switch between Host Antiviral and Antibacterial Responses. Mol Cell 71(4), 637–648 e635. doi: 10.1016/j.molcel.2018.07.010.

Perez, B., Mechinaud, F., Galambrun, C., Ben Romdhane, N., Isidor, B., Philip, N., et al. (2010). Germline mutations of the CBL gene define a new genetic syndrome with predisposition to juvenile myelomonocytic leukaemia. J Med Genet 47(10), 686–691. doi: 10.1136/jmg.2010.076836.

Peters, W., Cyster, J.G., Mack, M., Schlondorff, D., Wolf, A.J., Ernst, J.D., et al. (2004). CCR2-dependent trafficking of F4/80dim macrophages and CD11cdim/intermediate dendritic cells is crucial for T cell recruitment to lungs infected with Mycobacterium tuberculosis. J Immunol 172(12), 7647–7653. doi: 10.4049/jimmunol.172.12.7647.

Peters, W., Scott, H.M., Chambers, H.F., Flynn, J.L., Charo, I.F., and Ernst, J.D. (2001). Chemokine receptor 2 serves an early and essential role in resistance to Mycobacterium tuberculosis. Proc Natl Acad Sci U S A 98(14), 7958–7963. doi: 10.1073/pnas.131207398.

Pfeffer, P.E., Hopkins, S., Cropley, I., Lowe, D.M., and Lipman, M. (2017). An association between pulmonary Mycobacterium avium-intracellulare complex infections and biomarkers of Th2-type inflammation. Respir Res 18(1), 93. doi: 10.1186/s12931-017-0579-9.

Prevots, D.R., Loddenkemper, R., Sotgiu, G., and Migliori, G.B. (2017). Nontuberculous mycobacterial pulmonary disease: an increasing burden with substantial costs. Eur Respir J 49(4). doi: 10.1183/13993003.00374-2017.

Ramakrishnan, L. (2012). Revisiting the role of the granuloma in tuberculosis. Nat Rev Immunol 12(5), 352–366. doi: 10.1038/nri3211.

Ratnatunga, C.N., Lutzky, V.P., Kupz, A., Doolan, D.L., Reid, D.W., Field, M., et al. (2020). The Rise of Non-Tuberculosis Mycobacterial Lung Disease. Front Immunol 11, 303. doi: 10.3389/fimmu.2020.00303.

Ribechini, E., Eckert, I., Beilhack, A., Du Plessis, N., Walzl, G., Schleicher, U., et al. (2019). Heat-killed Mycobacterium tuberculosis prime-boost vaccination induces myeloid-derived suppressor cells with spleen dendritic cell-killing capability. JCI Insight 5. doi: 10.1172/jci.insight.128664.

Rivollier, A., He, J., Kole, A., Valatas, V., and Kelsall, B.L. (2012). Inflammation switches the differentiation program of Ly6Chi monocytes from antiinflammatory macrophages to inflammatory dendritic cells in the colon. J Exp Med 209(1), 139–155. doi: 10.1084/jem.20101387.

Roca, F.J., and Ramakrishnan, L. (2013). TNF dually mediates resistance and susceptibility to mycobacteria via mitochondrial reactive oxygen species. Cell 153(3), 521–534. doi: 10.1016/j.cell.2013.03.022.

Russell, D.G., Cardona, P.J., Kim, M.J., Allain, S., and Altare, F. (2009). Foamy macrophages and the progression of the human tuberculosis granuloma. Nat Immunol 10(9), 943–948. doi: 10.1038/ni.1781.

Ruth, M.M., and van Ingen, J. (2017). New insights in the treatment of nontuberculous mycobacterial pulmonary disease. Future Microbiol 12, 1109–1112. doi: 10.2217/fmb-2017-0144.

Sampath, P., Moideen, K., Ranganathan, U.D., and Bethunaickan, R. (2018). Monocyte Subsets: Phenotypes and Function in Tuberculosis Infection. Front Immunol 9, 1726. doi: 10.3389/fimmu.2018.01726.

Samstein, M., Schreiber, H.A., Leiner, I.M., Susac, B., Glickman, M.S., and Pamer, E.G. (2013). Essential yet limited role for CCR2(+) inflammatory monocytes during Mycobacterium tuberculosis-specific T cell priming. Elife 2, e01086. doi: 10.7554/eLife.01086.

Sanna, S., Pitzalis, M., Zoledziewska, M., Zara, I., Sidore, C., Murru, R., et al. (2010). Variants within the immunoregulatory CBLB gene are associated with multiple sclerosis. Nat Genet 42(6), 495–497. doi: 10.1038/ng.584.

Saunders, B.M., and Cooper, A.M. (2000). Restraining mycobacteria: role of granulomas in mycobacterial infections. Immunol Cell Biol 78(4), 334–341. doi: 10.1046/j.1440-1711.2000.00933.x.

Scott, H.M., and Flynn, J.L. (2002). Mycobacterium tuberculosis in chemokine receptor 2-deficient mice: influence of dose on disease progression. Infect Immun 70(11), 5946–5954. doi: 10.1128/iai.70.11.5946-5954.2002.

Serbina, N.V., Jia, T., Hohl, T.M., and Pamer, E.G. (2008). Monocyte-mediated defense against microbial pathogens. Annu Rev Immunol 26, 421–452. doi: 10.1146/annurev.immunol.26.021607.090326.

Shastri, M.D., Shukla, S.D., Chong, W.C., Dua, K., Peterson, G.M., Patel, R.P., et al. (2018). Role of Oxidative Stress in the Pathology and Management of Human Tuberculosis. Oxid Med Cell Longev 2018, 7695364. doi: 10.1155/2018/7695364.

Shi, C., and Pamer, E.G. (2011). Monocyte recruitment during infection and inflammation. Nat Rev Immunol 11(11), 762–774. doi: 10.1038/nri3070.

Silva Miranda, M., Breiman, A., Allain, S., Deknuydt, F., and Altare, F. (2012). The tuberculous granuloma: an unsuccessful host defence mechanism providing a safety shelter for the bacteria? Clin Dev Immunol 2012, 139127. doi: 10.1155/2012/139127.

Singh, T.P., Vieyra-Garcia, P.A., Wagner, K., Penninger, J., and Wolf, P. (2018). Cbl-b deficiency provides protection against UVB-induced skin damage by modulating inflammatory gene signature. Cell Death Dis 9(8), 835. doi: 10.1038/s41419-018-0858-5.

Spaulding, A.B., Lai, Y.L., Zelazny, A.M., Olivier, K.N., Kadri, S.S., Prevots, D.R., et al. (2017). Geographic Distribution of Nontuberculous Mycobacterial Species Identified among Clinical Isolates in the United States, 2009-2013. Ann Am Thorac Soc 14(11), 1655–1661. doi: 10.1513/AnnalsATS.201611-860OC.

Srivastava, S., Ernst, J.D., and Desvignes, L. (2014). Beyond macrophages: the diversity of mononuclear cells in tuberculosis. Immunol Rev 262(1), 179–192. doi: 10.1111/imr.12217.

Sturner, K.H., Borgmeyer, U., Schulze, C., Pless, O., and Martin, R. (2014). A multiple sclerosis-associated variant of CBLB links genetic risk with type I IFN function. J Immunol 193(9), 4439–4447. doi: 10.4049/jimmunol.1303077.

Swamydas, M., and Lionakis, M.S. (2013). Isolation, purification and labeling of mouse bone marrow neutrophils for functional studies and adoptive transfer experiments. J Vis Exp (77), e50586. doi: 10.3791/50586.

Tang, R., Langdon, W.Y., and Zhang, J. (2019). Regulation of immune responses by E3 ubiquitin ligase Cbl-b. Cell Immunol 340, 103878. doi: 10.1016/j.cellimm.2018.11.002.

Traynor, T.R., Kuziel, W.A., Toews, G.B., and Huffnagle, G.B. (2000). CCR2 expression determines T1 versus T2 polarization during pulmonary Cryptococcus neoformans infection. J Immunol 164(4), 2021–2027. doi: 10.4049/jimmunol.164.4.2021.

Venkatasubramanian, S., Cheekatla, S., Paidipally, P., Tripathi, D., Welch, E., Tvinnereim, A.R., et al. (2017). IL-21-dependent expansion of memory-like NK cells enhances protective immune responses against Mycobacterium tuberculosis. Mucosal Immunol 10(4), 1031–1042. doi: 10.1038/mi.2016.105.

Verma, D., Stapleton, M., Gadwa, J., Vongtongsalee, K., Schenkel, A.R., Chan, E.D., et al. (2019). Mycobacterium avium Infection in a C3HeB/FeJ Mouse Model. Front Microbiol 10, 693. doi: 10.3389/fmicb.2019.00693.

Wallner, S., Lutz-Nicoladoni, C., Tripp, C.H., Gastl, G., Baier, G., Penninger, J.M., et al. (2013). The role of the e3 ligase cbl-B in murine dendritic cells. PLoS One 8(6), e65178. doi: 10.1371/journal.pone.0065178.

Winthrop, K.L., Marras, T.K., Adjemian, J., Zhang, H., Wang, P., and Zhang, Q. (2019). Incidence and Prevalence of Nontuberculous Mycobacterial Lung Disease in a Large United States Managed Care Health Plan, 2008-2015. Ann Am Thorac Soc. doi: 10.1513/AnnalsATS.201804-236OC.

Wirnsberger, G., Zwolanek, F., Asaoka, T., Kozieradzki, I., Tortola, L., Wimmer, R.A., et al. (2016). Inhibition of CBLB protects from lethal Candida albicans sepsis. Nat Med 22(8), 915–923. doi: 10.1038/nm.4134.

Wolach, B., Gavrieli, R., de Boer, M., van Leeuwen, K., Berger-Achituv, S., Stauber, T., et al. (2017). Chronic granulomatous disease: Clinical, functional, molecular, and genetic studies. The Israeli experience with 84 patients. Am J Hematol 92(1), 28–36. doi: 10.1002/ajh.24573.

Xiao, Y., Tang, J., Guo, H., Zhao, Y., Tang, R., Ouyang, S., et al. (2016). Targeting CBLB as a potential therapeutic approach for disseminated candidiasis. Nat Med 22(8), 906–914. doi: 10.1038/nm.4141.

Yasuda, T., Tezuka, T., Maeda, A., Inazu, T., Yamanashi, Y., Gu, H., et al. (2002). Cbl-b positively regulates Btk-mediated activation of phospholipase C-gamma2 in B cells. J Exp Med 196(1), 51–63. doi: 10.1084/jem.20020068.

Zhu, L.L., Luo, T.M., Xu, X., Guo, Y.H., Zhao, X.Q., Wang, T.T., et al. (2016). E3 ubiquitin ligase Cbl-b negatively regulates C-type lectin receptor-mediated antifungal innate immunity. J Exp Med 213(8), 1555–1570. doi: 10.1084/jem.20151932.

