## Supplemental Data for "Decoding the role of CBLB for innate immune responses regulating systemic dissemination during Non-Tuberculous Mycobacteria infection"

**SUPPLEMENTARY FIGURE LEGENDS**

**Supplementary Figure 1**: *Cblb*-deficiency promotes NTM dissemination into the brain. *Six to eight*-wk-old OT-I-Tg (*Cblb*$^{+/+}$) and OT-I-Tg-Cblb$^{-/-}$ (*Cblb*$^{-/-}$) mice were infected either intravenously (I.V.; **A**) or intratracheally (I.T.; **B**) as described in Fig. 2. At indicated week PI, brain tissues were harvested and bacterial loads were quantified. Scatter plots with circles depict the individual mouse data, and horizontal line is the mean of the group. N=3-6 mice/group. IV infection data is representative of two independent experiments. **p≤0.01.

**Supplementary Figure 2**: CD11b$^{-ve}$ NK cell responses in the absence of *Cblb*. *Cblb*$^{+/+}$and *Cblb*$^{-/-}$ mice were infected as described in Fig. 2. At indicated week PI, tissues were harvested, single-cell suspensions were stained for NK cells (CD90$^{-}$, CD11b$^{+/-}$ NK1.1$^{+}$) using fluorochrome-conjugated antibodies, and analyzed by flow cytometry. **A & B.** Bar diagrams show the frequencies of CD11b$^{-}$ NK cells following IV and IT infections, respectively. IV infection data is representative of 2-3 independent experiments. Values are mean ± SD percent cells. N=3-6 mice/group. *p≤0.05 and **p≤0.01.

**Supplementary Figure 3**: Activated Neutrophils during NTM infection under *Cblb*-deficiency. *Cblb*$^{+/+}$and *Cblb*$^{-/-}$ mice were infected and cells were analyzed by flow cytometry as described in Fig. 8. Bar diagrams show MFI of CD44 on neutrophils in the lung (**A**) and spleens (**B**) infected by I.V. or I.T. route, respectively. Values are mean ± SD. N=3-6 mice/group. IV infection data is representative of 2-3 independent experiments. *p≤0.05, **p≤0.01, and ****p≤0.0001.

**Supplementary Figure 4**: CCL2 expression. Splenocytes or BMM were infected with 5 MOI of MAV104. Cells were harvested after 24hr PI, and subjected to qPCR analysis. Bar diagrams show fold change in gene transcript Vs. Uninfected *Cblb*$^{+/+}$. Data is pooled from two independent experiments. Values are mean ± SD. N=7 replicates/group. **p≤0.01.

**Supplementary Figure 1.**

**A.**

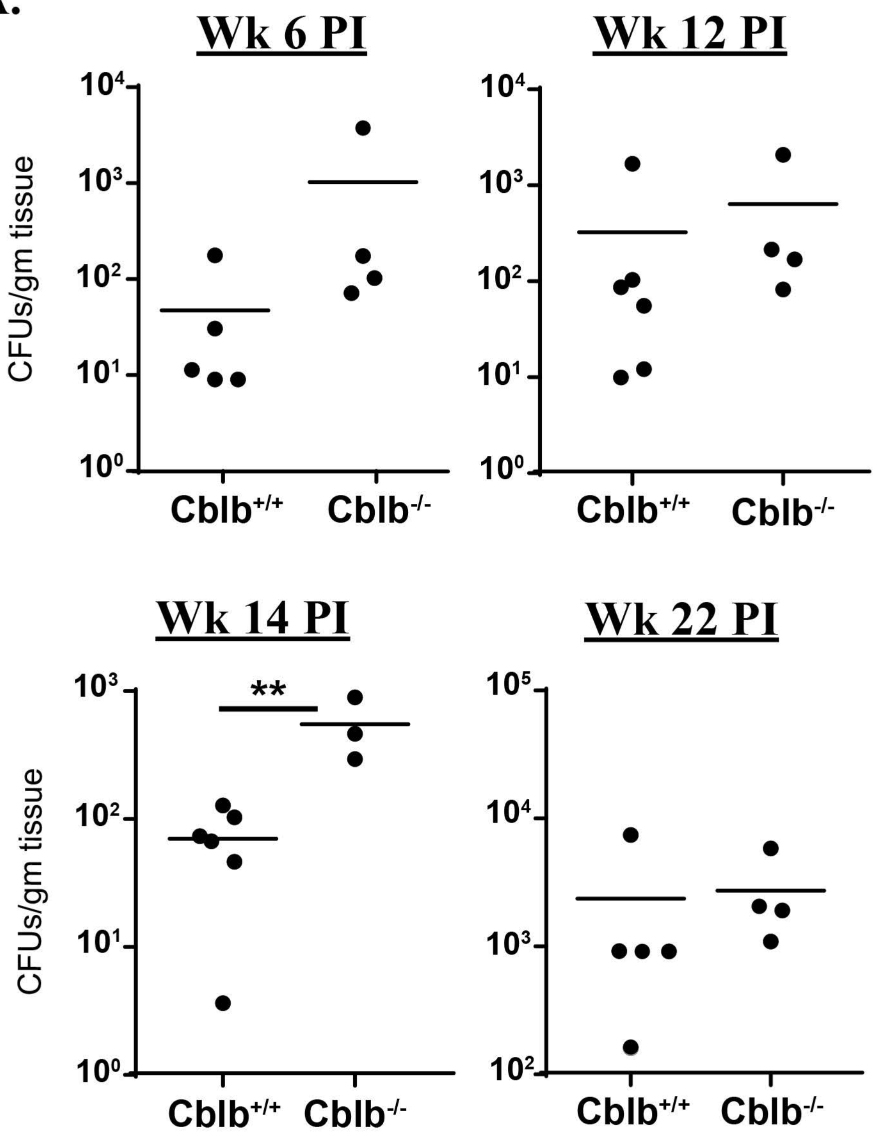


**Brain** **I.V.**

**B.**

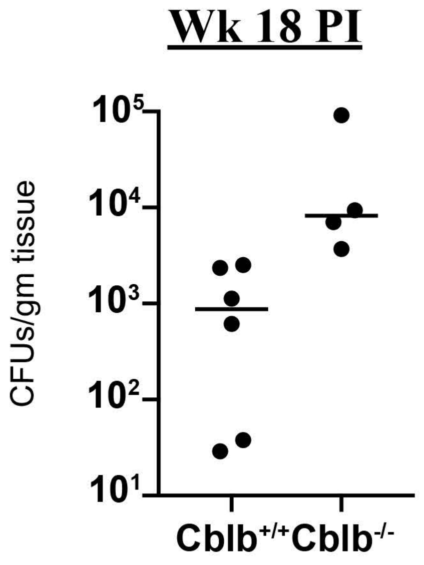


**Brain** **I.T.**

**Supplementary Figure 2.**

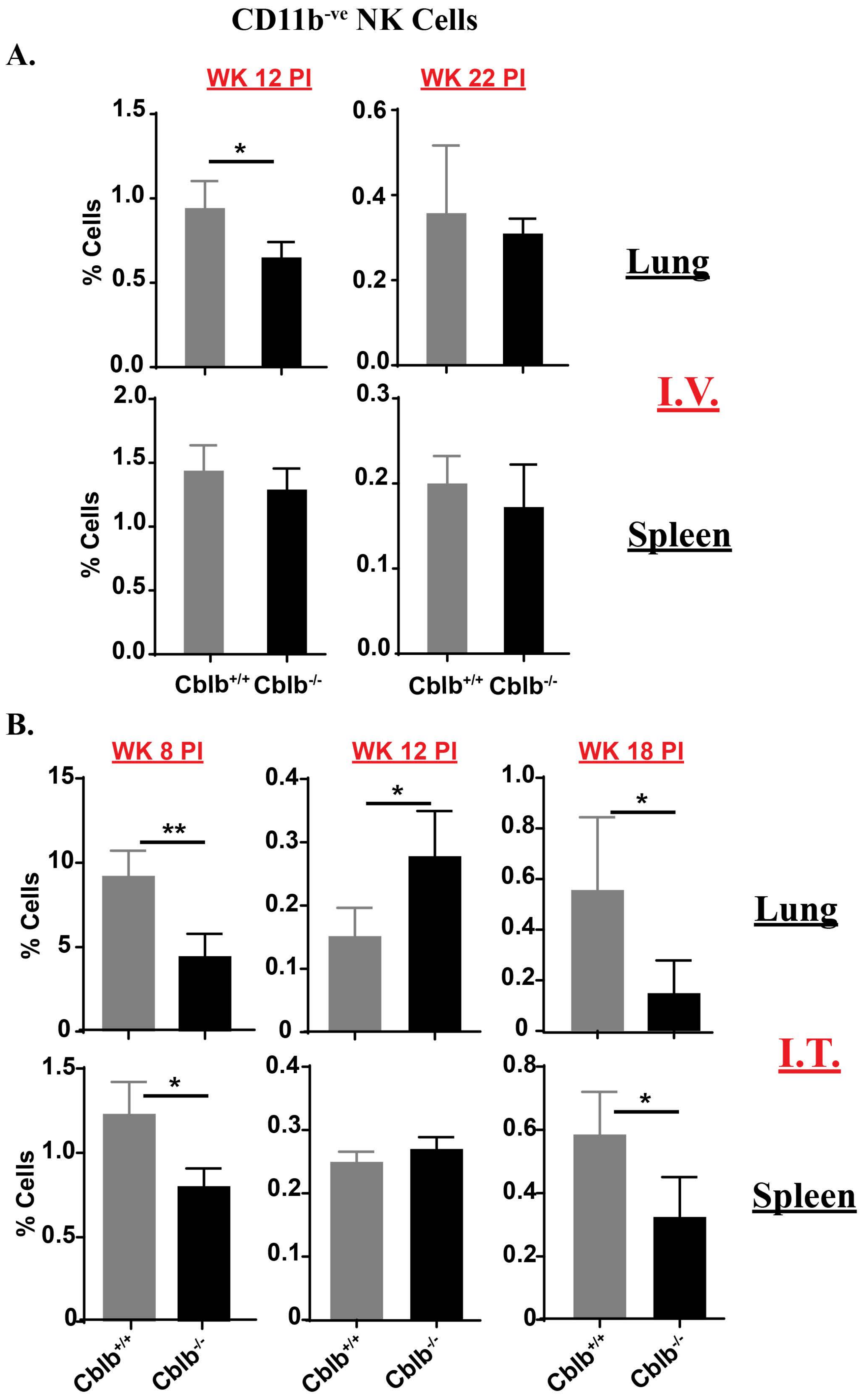

**Supplementary Figure 3.**

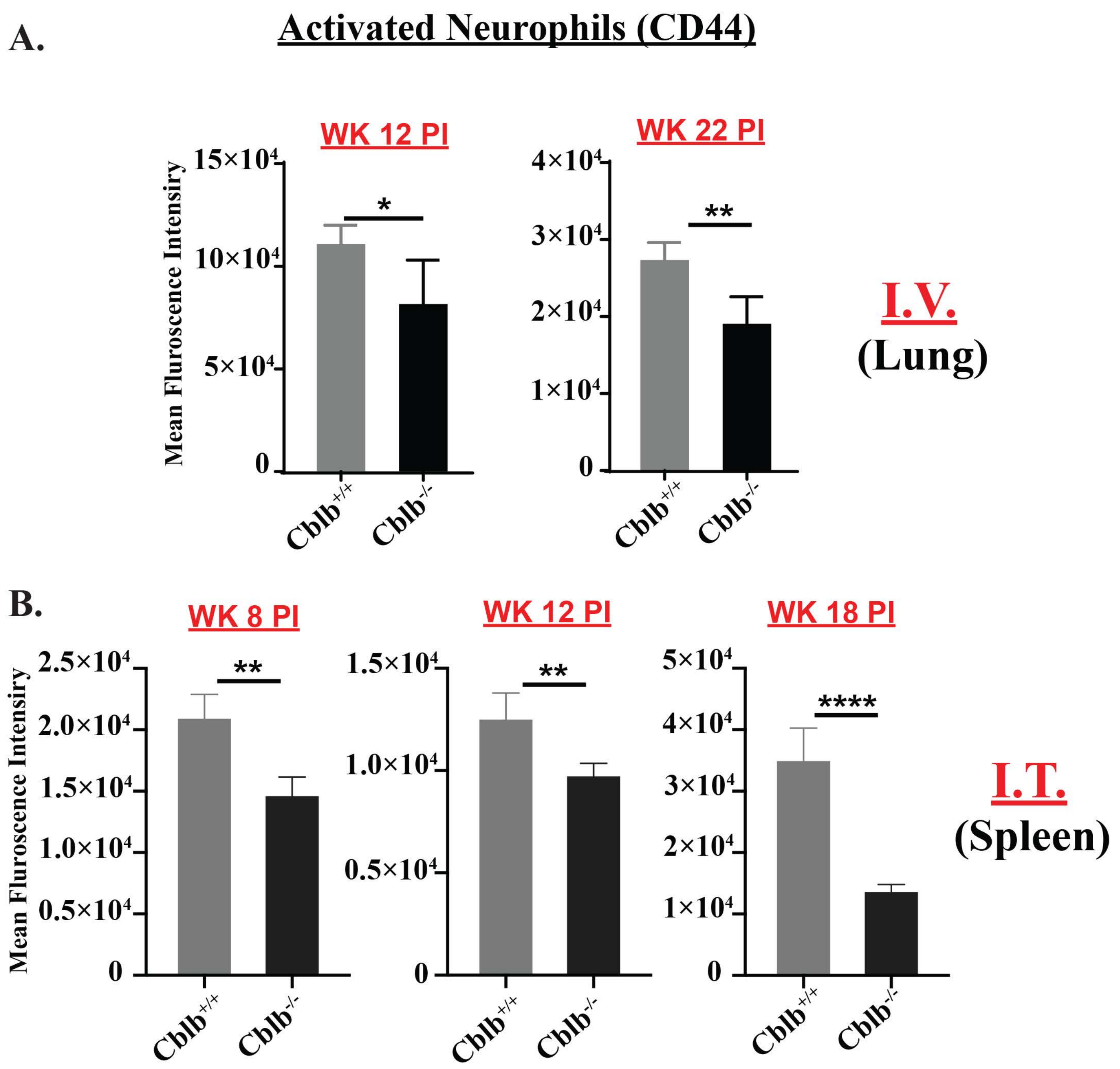

### Supplementary Figure 4.

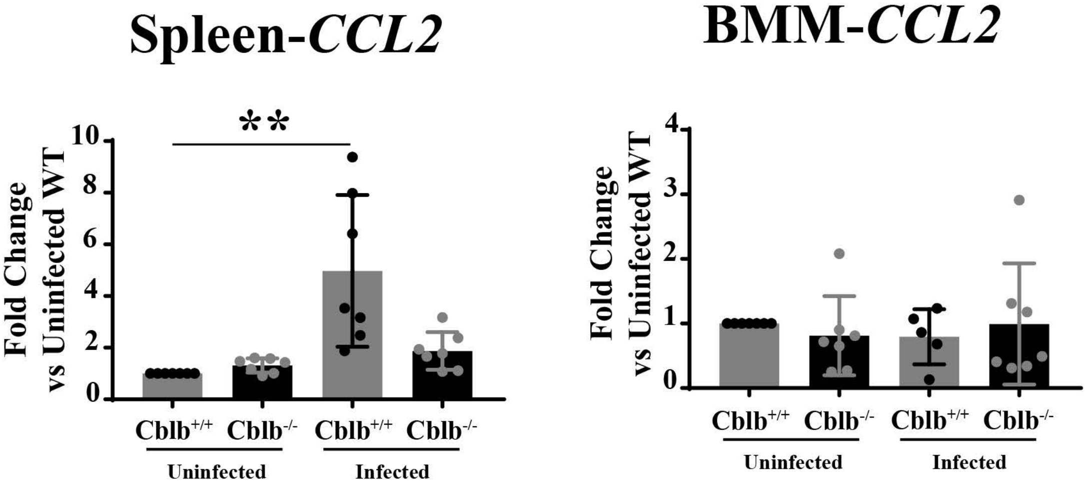
